# High-Density Multi-Distance fNIRS Enhances Detection of Brain Activity during a Word-Color Stroop Task

**DOI:** 10.1101/2025.03.12.642917

**Authors:** Jessica E. Anderson, Laura Carlton, Sreekanth Kura, Walker J. O’Brien, De’Ja Rogers, Parisa Rahimi, Parya Y. Farzam, Muhammad H. Zaman, David A. Boas, Meryem A. Yücel

## Abstract

**Significance:** Functional Near-Infrared Spectroscopy (fNIRS) enables neuroimaging in scenarios where other modalities are less suitable, such as during motion tasks or in low-resource environments. Sparse fNIRS arrays with 30mm channel spacing are widely used but have limited spatial resolution. High-density (HD) arrays with overlapping, multi-distance channels improve sensitivity and localization but increase costs and setup times. A statistical comparison of HD and sparse arrays is needed for evaluating the benefits and trade-offs of HD arrays.

**Aim:** This study provides a statistical comparison of HD and sparse fNIRS performance to inform array selection in future research.

**Approach:** We measured prefrontal cortex (PFC) activation during congruent and incongruent Word-Color Stroop (WCS) tasks using both Sparse and HD arrays for 17 healthy adult participants, comparing dorsolateral PFC channel and image results at the group level.

**Results:** While both arrays detected activation in channel space during incongruent WCS, channel and image space results demonstrated superior localization and sensitivity with the HD array for all WCS.

**Conclusions:** Sparse channel data may suitably detect activation from cognitively demanding tasks, like incongruent WCS. However, the HD array outperformed Sparse in detecting and localizing brain activity in image space, particularly during lower cognitive load tasks, making them more suitable for neuroimaging applications.

## 1 INTRODUCTION

The use of fNIRS has grown exponentially since its inception^1,2^ expanding our understanding of brain activity in a variety of contexts, such as in psychiatry^3–7^, naturalistic environments^8^, developmental research^9–11^, low-resource contexts and global health^11–17^, and for hyperscanning studies^18–21^. Benefits of functional Near Infrared Spectroscopy (fNIRS) over other neuroimaging modalities are well-defined for its comparative motion artifact resistance, temporal resolution, portability and wearability, operational requirements and ease of use, comfort, and general ecological validity^22^.

The traditional optode layout employed by most commercially-available fNIRS systems and studies arranges sources and detectors in a non-overlapping, 30-mm, grid pattern. This low-density or sparse arrangement suffers limited spatial resolution, sensitivity, and localization. For example, the ability to differentiate regions of activation is hindered because the channel density and spatial arrangement are directly related to the anatomical specificity^23^. The downstream impacts of this vary. One possible scenario is that the channels miss a region entirely; another is that channels may read signal from multiple nearby regions, and without enough spatial diversity of channels to specify regional activation, the activation from multiple regions essentially get averaged together. Either scenario prevents robust interpretation of brain behavior whether in magnitude, region specificity or both.

Additionally, Sparse arrays are known to exhibit poor fNIRS signal reproducibility because of non-uniform spatial sensitivity^24^. Though several works demonstrate that additional use of short-separation channels to the traditional Sparse array improves data quality and sensitivity to cerebral hemodynamics by enabling regression of hemodynamics from superficial tissue from the long channel measurements^25–27^, truly improved depth sensitivity additionally requires overlapping, high-density, multi-distance channels at lengths which allow for cortical sensitivity. This type of layout can improve spatial resolution as well as related signal characteristics such as partial volume blurring, spatial and depth sensitivity, localization of brain response, and inter-subject consistency^28^. These improvements are necessary to overcome limits of fNIRS’ ability to compare task-evoked response between conditions or between brain regions^29,30^, for example, and in applications toward current important work in malnutrition in global contexts^31^, brain disorders^32,33^, surgery assistance^34,35^, and brain-computer interfaces^36^ among others.

Recent advances in miniaturization of hardware components and fNIRS technology have made it possible to develop systems with high-density and overlapping, multi-distance channels. This approach, High-Density Diffuse Optical Tomography (HD-DOT), was first introduced in 2007^37^ and the first fiberless HD-DOT was introduced in 2016^38^. The fiberless nature makes a system ‘wearable’ and is important for approaching true ecological validity. HD-DOT attains high degrees of sensitivity and in fact, Eggebrecht *et al*. showed their system’s sensitivity approaches that of fMRI^39^.

It has been demonstrated that localization can be improved through various methods to resolve high-density fNIRS signal depth^31,40–43^. Currently, there are few commercially-available HD-DOT systems^44–46^ as well as several systems developed in labs^38,44,47,48^.

As more fNIRS and DOT options become available, it will be necessary for users to be able to quantitatively compare systems in order to select an optimal setup for the needs of their application. A key drawback of using an HD system over a Sparse system for a given regional coverage is the increased cost of resources such as materials – optodes, optode-to-cap attachments, control boards, other hardware – and computing processing required to efficiently process the larger amounts of data especially when considering whole-head arrays. When considering application to resource-limited settings, for example, the priority of this characteristic increases. Additionally, regardless of application, increasing the number and density of optode modules needed for the HD system especially in regions with hair incurs an increased setup and signal optimization time. The potential user needs to assess if the improved signal quality characteristics afforded by HD-DOT over traditional Sparse fNIRS are necessary to achieve the goals set out in their investigation and justify the disadvantages that come from having a denser array. It may be that the goal is to simply capture activation in a broad field-of-view, or to compare task evoked-response, localize activation, perform connectivity analysis, or achieve localization consistency, all of which may have varying degrees of improvement from a HD array. It is critical, therefore, that the field provides direct quantitative comparison of HD performance to that of traditional fNIRS arrays.

A relevant study by Shin *et al.* compared various fNIRS channel length combinations ranging from 15mm to 35mm and effectively compared HD to Sparse fNIRS. The signal comparison was evaluated by classification accuracy, which demonstrated that the combination of multi-distance and overlapping channels resulted in better classification accuracy than did a standard 30mm grid layout^40^. While this supports moving toward overlapping HD arrays for specific application, there remains need to statistically compare other metrics of fNIRS measurements of functional activity.

To that end, to the best of our knowledge there are two studies, from Fishell *et al*.^31^ and Frijia *et al.*^49^, which performed a direct comparison between a Sparse and HD layout with matching field-of-views and concluded HD arrays provide better localization of functional activity. The studies were in developing age populations and the array comparisons were made between their entire HD probe and a subset of channels to form a sparse array from the same data collection.

Fishell *et al.*’s fiber-based HD system was composed of three channel lengths (13mm, 29mm, 39mm) and spanned the temporal and occipital regions; the “Sparse” array measurements were compiled from the 29mm channel data. The image results qualitatively showed that the HD-DOT layout provides greatly improved localization of functional activity at the subject and group level and demonstrated inter-subject localization consistency, though the results did not report a statistical comparison between the array results. Additionally, the Sparse sub-layout did not include short-separation channels for superficial tissue regression and was not a grid pattern, so it therefore did not best represent what is currently most commonly seen in sparse fNIRS systems.

Frijia *et al.*’s study used the wearable, fiberless GowerLabs’ Lumo components for their HD layout with multiple channel lengths (ranging from 10mm to 45mm) primarily over the superior temporal lobes; the “Sparse” array uses only the data from a subset of non-overlapping channels whose length is between 20mm-24.5mm. Their HD results showed higher amplitude of hemodynamic response (HRF) compared to Sparse results in both channel and image space. Additionally, the SNR of their channel HRF was reduced in HD measurements compared to Sparse, and qualitatively the images showed improved and more consistent localization by the HD array. While these results take the comparison of sparse and HD arrays a step further, neither array included short-separation channels for regression of superficial tissue hemodynamics. Statistical comparison between array measurements was not provided.

The agreement among these studies’ findings is encouraging and reinforces the potential for improved localization with HD fNIRS. Building on this foundation, our study provides complementary evidence by conducting a direct statistical comparison of Sparse and HD fNIRS arrays. Furthermore, it extends prior work by examining array performance across different conditions, including varying cognitive loads, as well as applying short-separation regression, offering deeper insights into optimal array design for specific applications.

Our work performs a direct comparison of measured Word-Color Stroop (WCS) induced frontotemporal activation in 17 healthy adult subjects as detected by both a traditional grid-pattern, Sparse fNIRS array and a hexagonal-pattern, overlapping and multi-distance, high-density (HD) array to quantify the expected improvements afforded by the HD array. The WCS paradigm has been shown to elicit activation in the dorsolateral prefrontal cortex (dlPFC) due to the required response inhibition processing^50–57^ so our probes are designed to extend across the PFC. We have modeled a Sparse optode layout after one of the most commonly used commercially-available systems, ETG-4000 (Hitachi Medical Co., Tokyo, Japan) and our novel HD array is patterned after the NinjaNIRS^47^ layout and designed to have a field-of-view matching that of the Sparse array. After standard signal pre-processing and image reconstruction we generate concentration amplitude and t-statistics per subject for oxygenated and deoxygenated hemoglobin across WCS blocks for the channel data as well as brain surface vertices in image data. These metrics are both visualized and statistically compared at the group level. We find, in agreement with previous studies, that the HD array provides better localization than the Sparse array. We also find that the HD array captures, on average, stronger signal than does the Sparse array as defined by t-statistic per channel. The Sparse array is suitable for detecting, though not localizing, presence of activity for the incongruent WCS but not for congruent WCS. Our resulting comparison of layout and paradigm conditions provides a useful reference for future fNIRS users to make an evidence-based evaluation of optimal probe design for their application given experimental needs and limitations.

## 2 MATERIALS AND METHODS

### 2.1 Participants

Twenty-three healthy adult subjects were recruited from the Boston, MA area for this study via methods approved by the Boston University Charles River Campus Institutional Review Board (IRB Protocol 4502). Subjects gave written consent prior to the start of data collection. Subjects were not eligible for the study if they were outside the range of 18-89 years of age, had history of neurological trauma or neurological or psychiatric disorders, were currently taking psychoactive medications, were wearing a pacemaker or implantable cardioverter defibrillator, were wearing a deep brain stimulator, were wearing cochlear implants, had an uncorrected visual problem, or had a history of hearing problems. After subject exclusions due to technical errors (i.e. loss of signal transmission between devices during data collection) and poor data quality assessed during processing, 17 subjects remained whose data was used in analysis. Their demographics are as follows: mean age = 25.8 ± 4.3 years; 8 females, 9 males; 14 right-handed, 2 left-handed, 1 not reported; 10 White, 8 Asian, 1 Not Reported.

### 2.2 fNIRS Measurements

#### 2.2.1 Optode Arrays

We designed two optode layouts using the open-source AtlasViewer platform^58^. They are openly accessible at openfnirs.org. We refer to our “Sparse” array as one which represents a traditional arrangement, depicted in the top row of Figure 1. Its channels are in a grid pattern with lengths of 30mm. The centered and bottom-most optode is anchored to FPz and the optodes at the left and right bottom corners are anchored to T7 and T8, respectively. This layout includes 17 sources and 16 detectors for 52 channels, with an additional 8 detectors at 8mm separation (for a total of 24 detectors and 60 channels). Our multi-distance, high-density probe layout, referred to as the “HD” array, is shown in the bottom row of Figure 1. The HD array was designed such that its field-of-view is as close as possible to that of the Sparse array, given pattern allowance. The centered optode on the bottom row is similarly anchored to FPz; the left and right corner optodes are just outside of the T7 and T8 landmarks with spring lengths of 2mm to the landmarks. Following the hexagonal pattern recommended by von Lühmann *et al*., first nearest-neighbor (NN) channels are 8mm, second NN are 19mm, and third NN are 33mm. There are 25 sources and 58 detectors which produce 112 19mm channels and 94 33m channels, along with the additional 8 detectors at 8mm length short separation channels (for a total of 66 detectors and 214 channels). The files for each of these arrays are available to download at openfnirs.org/hardware/ninjaCap/.

**Figure 1:**
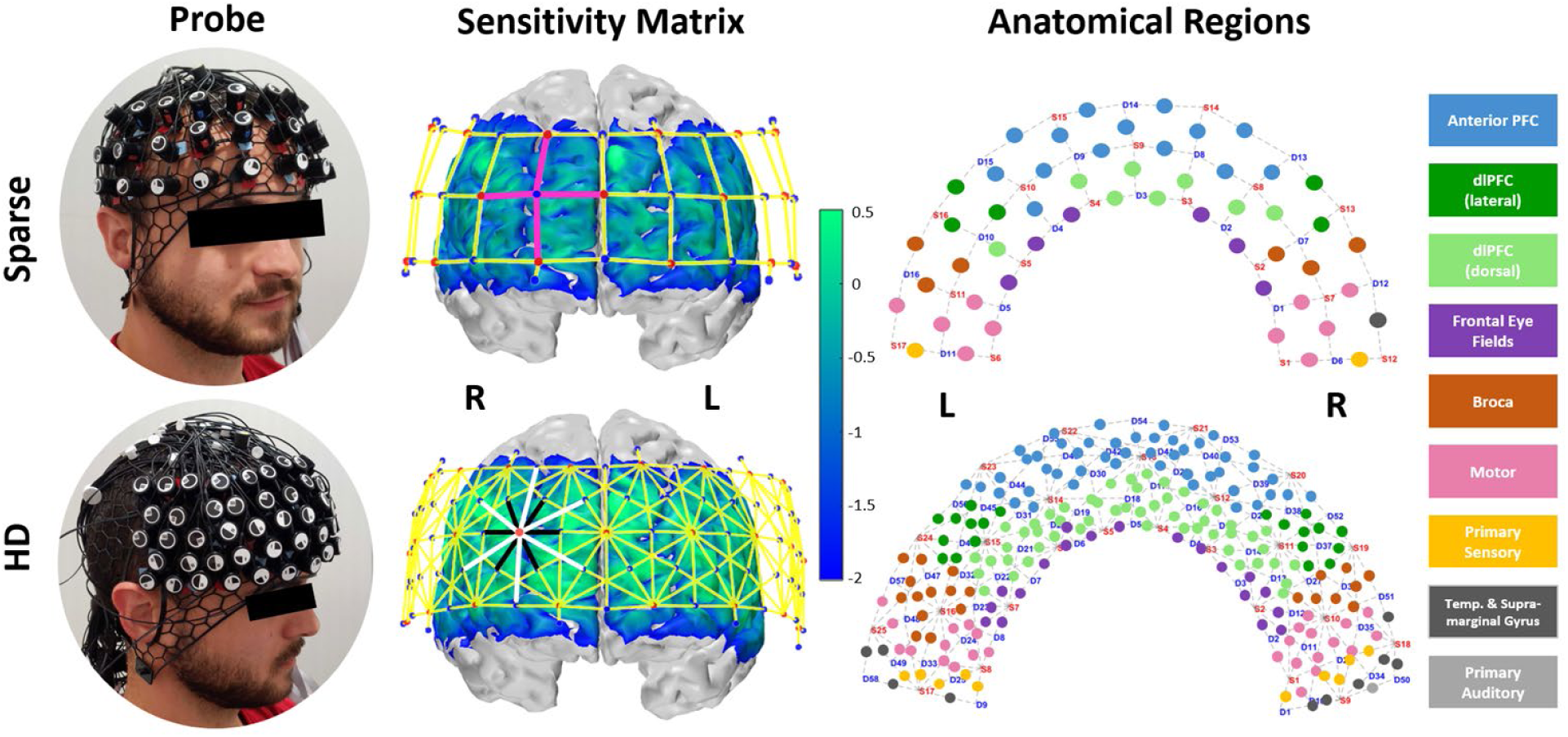
For each of the Sparse and High-Density (HD) probe design columns display physical appearance, sensitivity matrices via Monte-Carlo photon path modeling with probe overlay (red dots: sources, blue dots: detectors, pink lines: emphasize ‘grid’ layout of sparse array’s 30mm channels, black/white lines: emphasize ‘hexagonal’ layout of HD array’s 19mm/33mm channels), and Brodmann areas underlying each channel. Sensitivity profile is on a log 10 scale; vertices with values > 0.01 are not masked and not considered part of the relevant sensitivity profile.

To identify Brodman regions underlying each channel, we employed AtlasViewer’s “project channels to surface” function to calculate MNI coordinates for each channel of our Sparse and HD array. We then utilized the BioImage Suite Platform (www.bioimagesuite.org) which maps the Colin27 brain surface (as used in AtlasViewer and therefore in creating our digital probe design) to the Talairach atlas surface and identifies Brodman regions from the calculated MNI coordinates^59^. In our case, not every channel was assigned a region from these steps because the coordinate was not fully projected to the cortex. To complete the anatomical labeling of our probes as shown in the right-hand column of Figure 1, we developed four steps. The first is spatial interpolation per array, meaning that if an unlabeled channel is surrounded by channels with uniform labeling, that label was applied. Next, we compare an overlay of Sparse and HD channel labels; if one array’s unlabeled channel overlaps or is surrounded by labeled channels from the other array it was assigned correspondingly. Thirdly, within each array we assigned still unlabeled channels with that of the matching channel in the other hemisphere. We acknowledge no individual brain, and thus the Colin27 template itself, is perfectly symmetric across hemispheres but at the group level symmetry can be assumed for the sake of region identification. Finally, for remaining unlabeled channels, we projected them to the cortex by applying incremental coordinate adjustments in the BioImage platform.

#### 2.2.2 Hardware

For both the Sparse and HD layouts we used NIRSport2 optodes and systems (four cascaded NIRSport2 16×16 devices for HD and three for Sparse) and Aurora data acquisition software (NIRx, Berlin, Germany). We used the dual-tip sources, which emit light at 760nm and 850nm, and silicone photodiode detectors. The sampling frequencies were

24.4Hz for Sparse and 17.5Hz for HD. We manually optimized the spatial multiplexing of the HD array from its default settings in order to achieve its sampling frequency. We used fully customizable, conformable NinjaCap^60^, 3D printed in three sizes (54cm, 56cm, 58cm circumference) with NinjaFlex (Fenner Precision Polymers, Lititz, Pennsylvania, United States), a flexible thermoplastic polyurethane (TPU) filament. Each cap included a reference marker for the Cz landmark. The open webbing design of the cap improves accessibility to maneuver hair to improve scalp coupling across diverse demographics. This cap structure also allows more escape of heat, thus increasing subject comfort.

### 2.3 Experimental procedures and paradigm

#### 2.3.1 Experimental Procedure

For a given session, a subject first wore one array while performing the task, then after an approximately 30-minute break while optodes were moved to the other cap, the subject repeated the task while wearing the other array. We randomized the order in which the arrays are worn to ensure roughly half the data was from subjects who first wore the Sparse array and half from subjects who first wore the HD array.

To place the head caps, we first measured circumference, distance from inion (Iz) to nasion (Nz), and distance from ear-to-ear or LPA (T9) to RPA (T10). The cap for the selected measurement was placed to align both the Cz landmark and FPz optode on the cap to the subject’s head. Cz was measured as the intersection of the midpoint of a line connecting the Iz and Nz landmarks with the midpoint of a line connecting the LPA and RPA landmarks. FPz was measured from Iz as 10% of the total distance from Iz to Nz. Due to the nature of skull shape and size variety, cap placement according to one of these landmarks did not always converge to placement according to the other landmark. In cases where this was observed, alignment to FPz was prioritized because the array is focused on the frontal region. To reduce likelihood of hair falling between the optodes and scalp especially on the forehead and temple we first requested subjects look upward and set the frontal portion of the cap on first, then instructed them to look straight ahead and gently unfolded the cap over the rest of the head from front-to-back with one hand while lightly holding the front of the cap in place with the other. Scalp coupling optimization generally required more time and care for the HD array due to the increased number of optodes, which not only required more optodes be attended to, but additionally prevented easy displacing of hair from under one optode without accidentally pushing it under the next. Generally, moving hair was performed with a cotton-tipped applicator. To achieve best signal quality, with the permission of the subject we sometimes applied clear ultrasound gel via the applicator. The optode was removed from the grommet, gel applied to the region as hair was pushed to the sides, and optode replaced; the gel aided to maintain hair position so that it did not move back under the optode. While sometimes good scalp-coupling – as defined by the Aurora ‘green’ thresholds of covariance < 2.5% and raw signal detection > 3mV – was achieved near immediately for most or all channels especially with bald or short hair, we would spend a maximum of approximately 20 minutes to optimize the scalp-coupling before moving on. Room lighting was provided by halogen floor lamps rather than overhead fluorescent bulbs in order to avoid interference from ambient pulsed light. To prevent effects on frontal detectors from ambient signal emitted by the paradigm-presentation laptop, we placed a black shower cap on the subjects’ head over the optodes.

Duration of sessions from the time of consent to completion was approximately 2-2.5 hours.

#### 2.3.2 Paradigm

Our modified WCS task (Figure 2) was based on previous work by Jahani *et al.*^51^. We compiled and presented it via PsychoPy^61^ with the full paradigm available on GitHub (https://github.com/andersonjessie/WordColorStroop). Our WCS had two conditions: an easier congruent task, and more difficult incongruent task. A trial of the congruent condition would display two words on a black background. Prior to a run, instructions for the congruent condition were provided on the screen and read aloud to the subject as follows: “For the following, two words will appear. Press either the Left Arrow or Right Arrow to indicate which word matches its font color. Press space to proceed to the practice.” The subject practiced twice with feedback, with additional practice if needed for clearer understanding. The design of the congruent task ensured that per trial, one word was congruent to its font color and the other was incongruent to its font color. A trial of the incongruent condition would display four words on a black background. Instructions for incongruent were provided on the screen and read aloud to the subject as follows: “For the following, a word will appear in the middle and a set of three words will appear above it. Press the Left Arrow, Up Arrow, or Right Arrow to indicate which word of the top row (left, middle, and right, respectively) describes the font color of the word below. Press space to proceed to the practice.” Again, the subject practiced twice with feedback, with additional practice if needed for clearer understanding. Instructions and practices were repeated when the subject performed another run for the second array. The design of the incongruent task ensured all four words (the prompt word and three option words) were always incongruent to their own font color. In both conditions, the location of the correct response was randomized.

**Figure 2:**
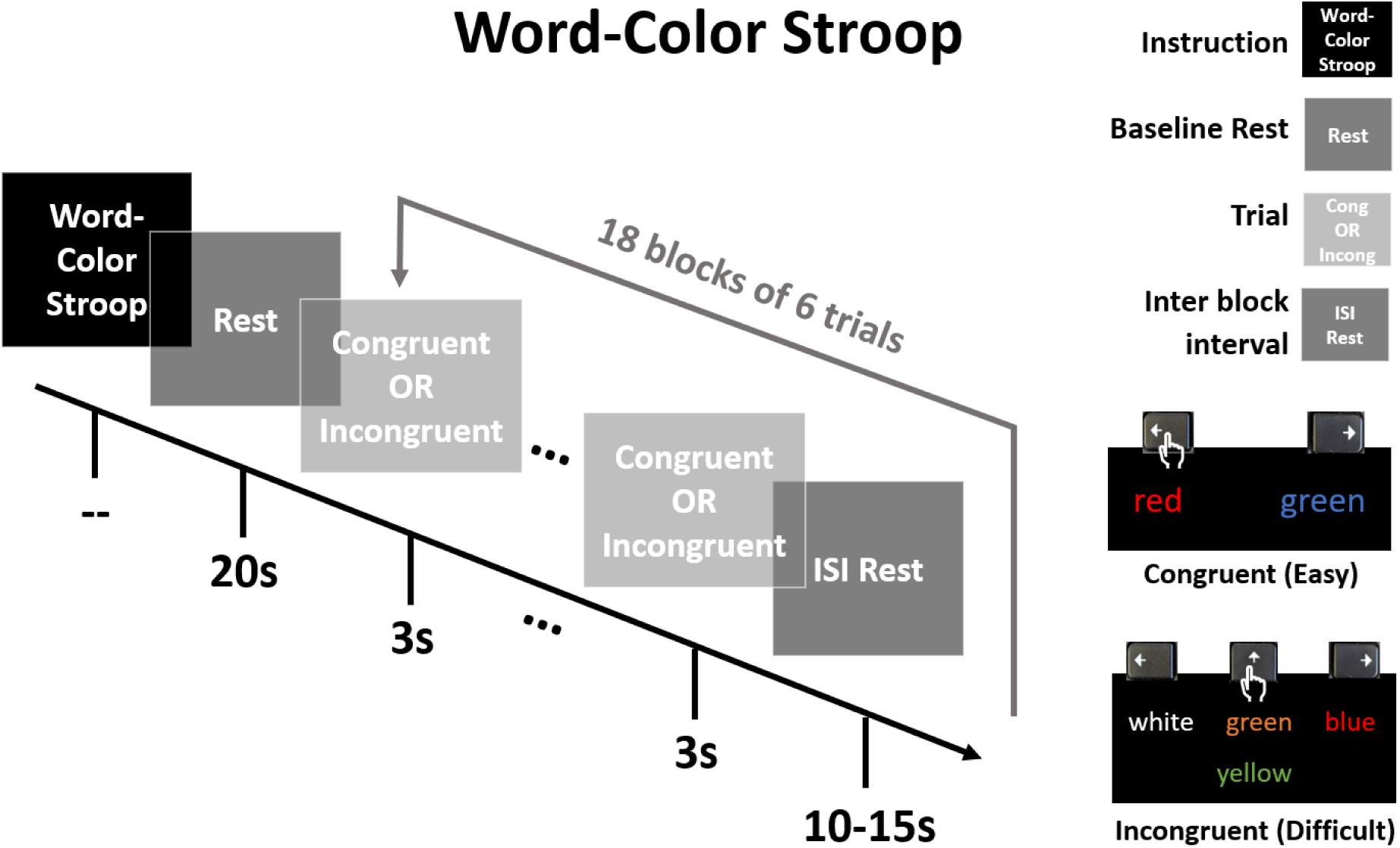
Word-Color Stroop paradigm adapted from Jahani, et al. After instruction and initial rest, 18 blocks of 6×3-s trials each were presented with a jittered inter-block interval (10-15 s). A given block consisted of either all congruent (Easy) trials, or all incongruent (Difficult) trials. The lower-right legend demonstrates accurate user keyboard press for each condition. Order of blocks was randomized for a total of 9 blocks of each condition. Total run time approximates 11min.

After completing instruction and practice, a run began with a 20 s rest then 18×18 s blocks with interstimulus rest jittered between 10-15 s. There were 9 blocks of each condition with total block order randomized. Each block had 6 trials with duration 3 s per trial, and a given block was either all congruent or all incongruent trials with no feedback on accuracy provided to the subject. Total run time was approximately 11 minutes.

### 2.4 Data Pre-Processing

We performed the following pre-processing steps in the open-source Homer3 platform^62^. First, channels with raw signal levels less than 0.001 intensity units (for NIRSport2 data, V) or with a signal-to-noise ratio value less than 5 were pruned from further analysis (hmrPruneChannels). Intensity was converted to optical density (OD) (hmrR_Intensity2OD). We used the SplineSG motion correction function which applies both spline-interpolation and Savitzky–Golay filtering to correct for baseline shifts and spikes, respectively (hmrR_MotionCorrectSplineSG)^63^, setting recommended parameter p value for smoothing to 0.99 and using a 10 second frame size. A low-pass filter (hmrR_BandassFilt: Bandpass_Filter_OpticalDensity) of 0.5 Hz was used to remove high-frequency noise such as cardiac signal before OD was converted to oxygenated-, deoxygenated-hemoglobin concentrations (HbO, HbR, HbT, respectively) via the Beer-Lambert Law (hmrR_OD2Conc)^64^. Upon visual inspection of raw signal, OD, and concentration timeseries data, it was apparent that some of the slow motion artifacts in the temporal regions could not be fully corrected via available detection or correction functions, so we performed manual rejection of blocks which contained such artifacts. We then applied a general linear model (hmrR_GLM) to estimate the hemodynamic response function (HRF) from two seconds prior to block onset to 23s after block onset. This captures 5s after stimulus offset, at 18s, in order to cover the recovery period. In the GLM we used ordinary least squares and applied a consecutive sequence of gaussian functions of 1 s width and step. For each long channel we use the short-separation (8mm) channel with greatest correlation to regress superficial tissue signal. The GLM included a polynomial drift correction of order 3.

If a subject had fewer than half the maximum blocks remaining in analysis due to manual block rejection (i.e. if they had four or fewer blocks for either condition) they were excluded from further analysis, which was the case for four subjects.

### 2.5 Image Reconstruction

To generate our sensitivity matrices per array, we performed forward modeling of photon migration via AtlasViewer’s “runMCXlab” function. This uses the MCXLAB package^65^ to perform a Monte Carlo simulation in which the random walk of photons between each source and detector is mapped to generate the sensitivity profiles^66–68^ ^66–68^. This represents the spatial sensitivity of each measurement channel to changes in cortical absorption. We chose to use the default parameters for photons launched (1e7) per optode and used wavelength-appropriate tissue properties for scalp, CSF, gray matter, and white matter tissues^27^. Each row of the sensitivity matrix represents the sensitivity profile of an individual channel. By summing over all channels, one can visualize the total sensitivity of the probe design as shown in the middle column of Figure 1.

For a point activation on a cortical surface element, localization error (shown in Supplementary material Figure S1) can be summarized as the Euclidian distance between the true center of activation and center of mass of the reconstructed image. The position of center of mass *r_CM_* is calculated as:

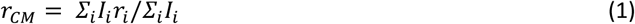

Where the *i*th surface element had intensity *I_i_* at position *r_i_*.

Resolution profiles *R_CM_*, also available in Supplementary material Figure S1, were calculated at each surface element by:

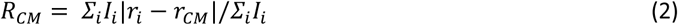

Image reconstruction was performed to project optical density in channel space to concentration on the cortical and scalp surfaces. To do this, the estimated HRF output for HbO and HbR from the GLM were first converted back to optical density. Channel space optical density measurements can be expressed as the product of the sensitivity matrix and the concentration changes on the cortical surface using the forward model:

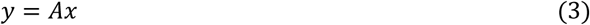

Where *y* is the channel space measurements in optical density, *A* is the sensitivity matrix and *A* is the changes in HbO and HbR on the cortical surface. To obtain these cortical measurements of HbO and HbR, we could solve the corresponding inverse problem:

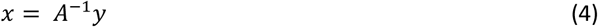

Reconstructing images on both the brain and scalp simultaneously has been shown to improve image resolution and localization^27,42^. Since the scalp is significantly more sensitive than the brain, spatially variant regularization is used to tune the reconstruction to the appropriate depth^42^. This is done by rescaling *A* as *A*^ using the diagonal matrix *L* defined as:

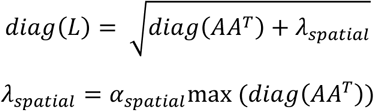

Then,

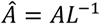

The parameter *α_spatial_* is used to control the reconstruction depth and is set to 0.001.

This inverse problem is ill-posed and underdetermined, however, so Tikhonov regularization was used to improve the estimation of *x*. The regularization parameter *α_meas_* was used to scale the measurement covariance such that, when combined with the spatially variant regularization, the inverse problem became:

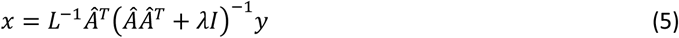

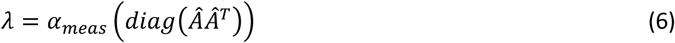

The parameter α_meas_ was used to smooth the image and was set to 0.001. We refer to this method as “brain and scalp” image reconstruction throughout.

To visualize a subject’s given set of vertices’ time course, we extracted the Homer GLM output HRF concentration data, generated the brain and scalp image for each second of HRF results, and plotted the mean of the vertices’ reconstructed image concentration over time.

### 2.6 Statistical Analysis

#### 2.6.1 Region of Interest

We selected the dorsolateral PFC (dlPFC) as our region of interest (ROI) as that is where we expect greatest activation to Stroop task based on previous studies^54^. We included channels with this labeling in Figure 1 and with similar field of view, excluding the ones assigned as dlPFC (dorsal) which fall in the medial area. In selecting ROI channels, we preserved symmetry of channels selected within and between each array, as well as account for inter-subject brain variability, which led to including several channels not initially marked as in the dlPFC. (Sparse ROI channels included: 8 dlPFC, 3 Anterior PFC, 1 Broca. HD ROI channels included: 42 dlPFC, 4 Anterior PFC, 2 Broca, 1 Frontal Eye Fields). The channels from each array selected per ROI appear in the left column of Figure 3. Per ROI, there are 6×30mm channels from the Sparse array, 11×19mm channels from the HD array, and 14×33mm channels from the HD array. In selecting image space ROIs, because both the Sparse and HD array have the same total vertices and placement we selected exactly matching ROI vertices. We first identified the vertices whose sensitivity to the HD ROI channels were above the same 0.01 threshold used in visualizing the sensitivity matrices. We then kept the vertices with sensitivity above the average per from each ROI.

**Figure 3:**
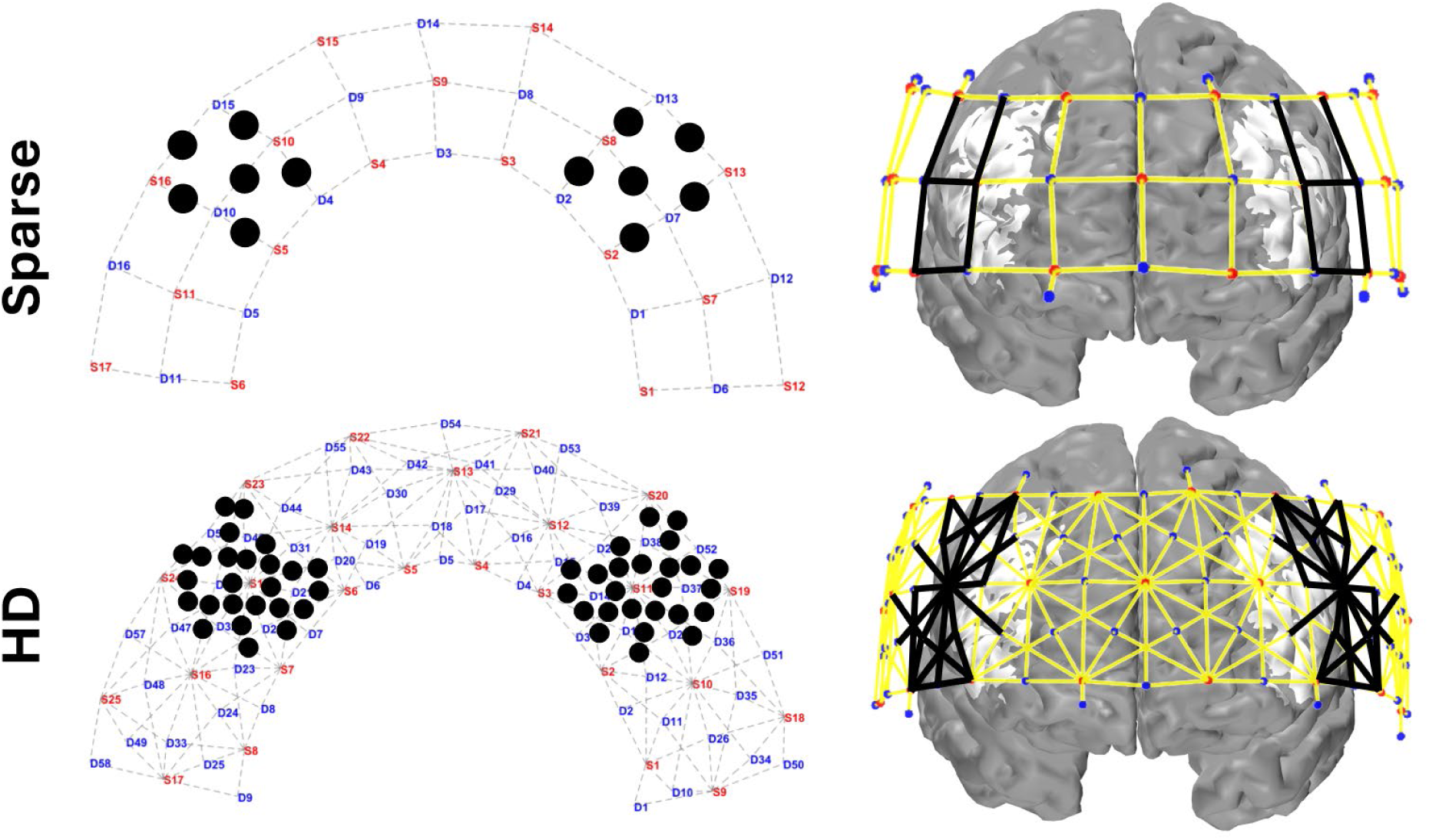
Channels and vertices selected in the region of interest for the Sparse and HD array. On the left panel, black dots mark the center of each channel included in the ROIs. On the right panel, black lines indicate the channels included in the ROIs and the white (unshaded) region of the brain indicates the vertices included. Vertices for both arrays are chosen based on those sensitive to the HD ROI channels.

#### 2.6.2 Array Comparison Statistics

We analyzed the GLM-processed concentration timeseries data for HbO and HbR, and for both congruent and incongruent conditions. For each block at a given channel of an individual subject, we calculated a delta concentration by subtracting the mean of the concentration time course from-2 to 0 seconds prior to block onset from the mean of the concentration time course between 7 to 18 seconds after the block onset. The mean delta concentration across blocks was divided by the standard error across blocks to produce a t-statistic particular to that concentration, WCS condition, channel, fNIRS array, and subject. For visualization purposes, group-level concentration means and standard error from across subjects’ own block-averages was used in calculating group-level t-statistics per channel.

To compare Sparse and HD HbO results, we selected from each subject the channel per left and right ROI which had the highest t-statistic. For HbR results, we selected based on lowest t-statistic with the evidenced expectation that HbR decreases when HbO increases for typical brain activation^69^. We performed a paired Student’s *t*-test to statistically compare array performance.

To analyze in image space, we first reconstructed the image of each block based on the peak minus baseline values per channel previously calculated. We then used similar methodology to analyze the image data as used to analyze channel data, applying it to the mean of the 25 image vertices with highest t-statistics.

## 3 RESULTS

### 3.1 Data Quality and Retention

Channel quality results are presented in *Table 1*. Here we focus on the report of all length HD channels. By paired Student’s *t*-test, there was no significant difference between the SNR of the Sparse Array channels and the SNR of the HD Array channels (p = 0.36). There was, however, a significant difference in the number and percentage of channels pruned from each cap (p = 4.3e^-7^ and p = 2.1e^-7^, respectively). The differences were also significant when looking specifically within the ROIs (p = 4.5e^-4^ and p = 8.7e^-4^ for number and percentage of channels pruned, respectively). The number of channels kept (pruned) in the analysis were, on average, 196.1 ± 9.4 (17.9 ± 9.4) HD channels and 59.6 ± 0.9 (0.4 ± 0.9) Sparse channels across the entire array; within our ROIs there were on average 48.3 ± 1.6 HD channels and 12.0 ± 0.1 Sparse channels included in the analysis. There was no significant difference between the number of blocks available for a given WCS task when the subject wore the Sparse array versus when they wore the HD array (p = 0.36 for incongruent, p = 0.21 for congruent by paired Student’s *t*-test). There was also no significant difference between the number of blocks available when wearing a given array for the incongruent versus congruent task (p = 0.31 for the Sparse array, p = 0.36 for the HD array via paired Student’s *t*-test).

**Table 1:** Channel signal quality metrics are provided in terms of mean and standard deviation across subjects. Congruent and incongruent WCS blocks are blocks remaining after rejections due to motion artifacts. * for p < 0.05, ** for p < 0.01, *** for p < 0.001 by paired t-test.

|  | SNR | Ch Pruned, total | Ch Pruned (%), total | Ch Pruned, ROI | Ch Pruned (%), ROI |
| --- | --- | --- | --- | --- | --- |
| **Sparse** | $54.9 \pm 17.0$ | $0.4 \pm 0.9$ | $0.6 \pm 1.4$ | $0.0 \pm 0.1$ | $0.2 \pm 1.0$ |
| **HD** | $51.6 \pm 15.9$ | $17.9 \pm 9.4$ | $8.4 \pm 4.4$ | $1.7 \pm 1.6$ | $3.5 \pm 3.2$ |
| **HD, 33mm** | $40.9 \pm 13.5$ | $14.8 \pm 7.3$ | $15.7 \pm 7.7$ | $1.5 \pm 1.4$ | $5.4 \pm 4.9$ |
| **HD, 19mm** | $57.3 \pm 18.6$ | $2.7 \pm 3.0$ | $2.4 \pm 2.7$ | $0.2 \pm 0.5$ | $1.1 \pm 2.1$ |

### 3.2 Performance Metrics

*Table 2* provides the group accuracy and response times (RT) per array and WCS condition of 16 subjects whose performance metrics were recorded and available for analysis. Between arrays, there was no significant difference in the accuracy or RT (p = 0.54 for congruent accuracy, p = 0.94 for incongruent accuracy, p = 0.40 for congruent RT, p = 0.99 for incongruent RT, via paired Student’s *t*-test). There was not a significant drop in accuracy when performing incongruent task as compared to congruent (p = 0.05 for Sparse array accuracy, p = 0.07 for HD array accuracy via paired Student’s *t*-test); however, there was a significant difference in response times between the conditions (p = 9.11e^-10^ for Sparse array RT, p = 4.24e^-7^ for HD array RT, via paired Student’s *t*-test).

**Table 2:** Performance metrics are provided as average across subjects’ trial averages. Three asterisks indicate p < 0.001.

|  | Accuracy (%) |  | Response Time (sec) |  |
| --- | --- | --- | --- | --- |
|  | Congruent | Incongruent | Incongruent | Congruent |
| **Sparse** | $97.8 \pm 4.7$ | $86.8 \pm 21.9$ | $1.57 \pm 0.24$ *** | $97.8 \pm 4.7$ |
| **HD** | $97.1 \pm 8.3$ | $87.2 \pm 18.5$ | $1.57 \pm 0.19$ *** | $97.1 \pm 8.3$ |

### 3.3 Functional Activation

For each WCS condition and array, group-averaged mean HbO and t-statistic are shown in channel space in Figure 4. The reconstructed image results for group-averaged HbO mean and t-statistics are shown in Figure 5. Functional activation was present in the lateral regions for the HD array for both conditions. Functional deactivation was also present in the medial PFC for the HD array, more strongly during incongruent WCS than during congruent WCS. The pattern of activation in the Sparse image is difficult to distinguish and for the most part opposite of what is expected according to the channel group results. Due to variation of channels pruned per subject, not all the group channel calculations had the same number of subjects whose data are contributing. On average, HD channels’ group data had 15.6 ± 2.4 subjects per channel and Sparse channels’ group data had 16.9 ± 0.4 subjects per channel. For visual and comparative continuity across caps and between channel and image space, the maximum *n* of 17 (with two-tailed α=0.05) was used to calculate the critical t-statistic value of 2.1 for group data regardless. This number is used here only for visual thresholding of group data.

**Figure 4:**
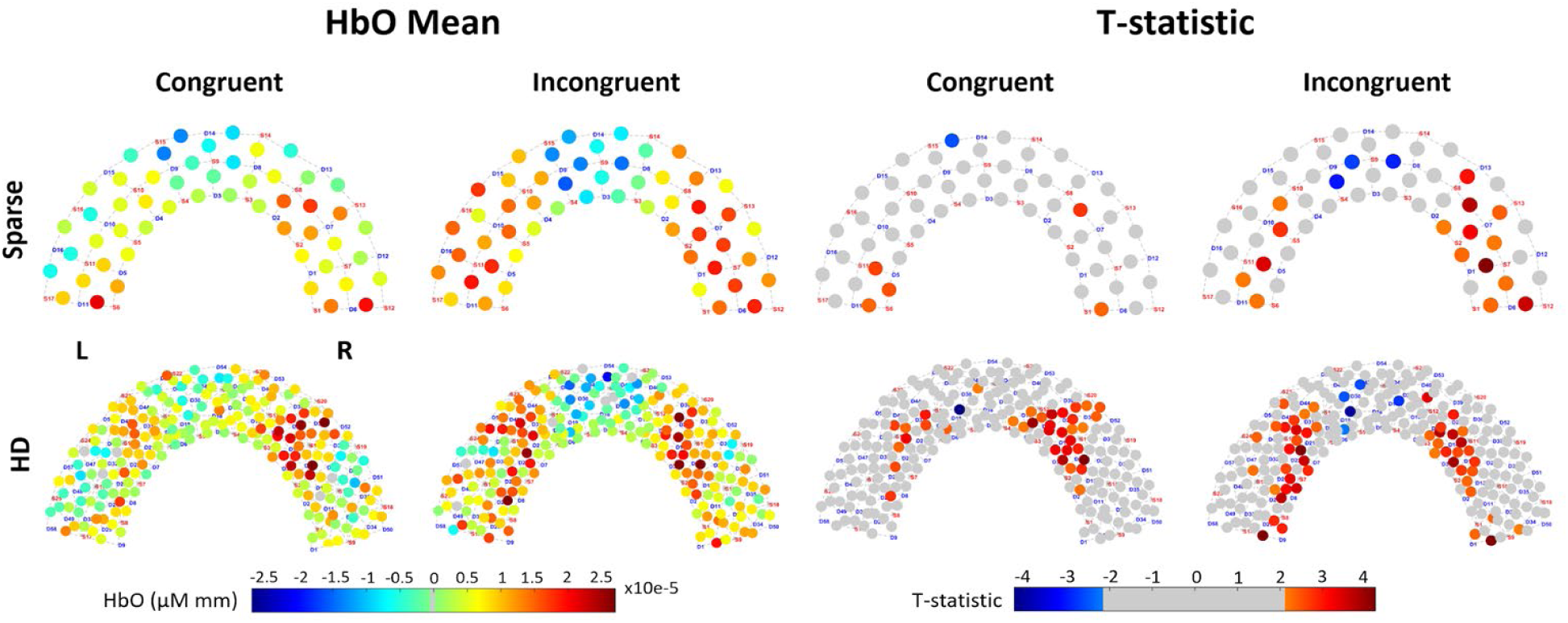
Channel space brain response recorded by Sparse and HD arrays during WCS, from Superior view. “HbO Mean”: Group-average hemodynamic response (HbO) for each channel, averaged across 7 to 18 seconds of the blocks for each condition. “T-statistic”: Group-averaged t-statistic of each channel is plotted. Color-scale is grey for absolute values less than t-crit = 2.12 as calculated for 17 subjects.

**Figure 5:**
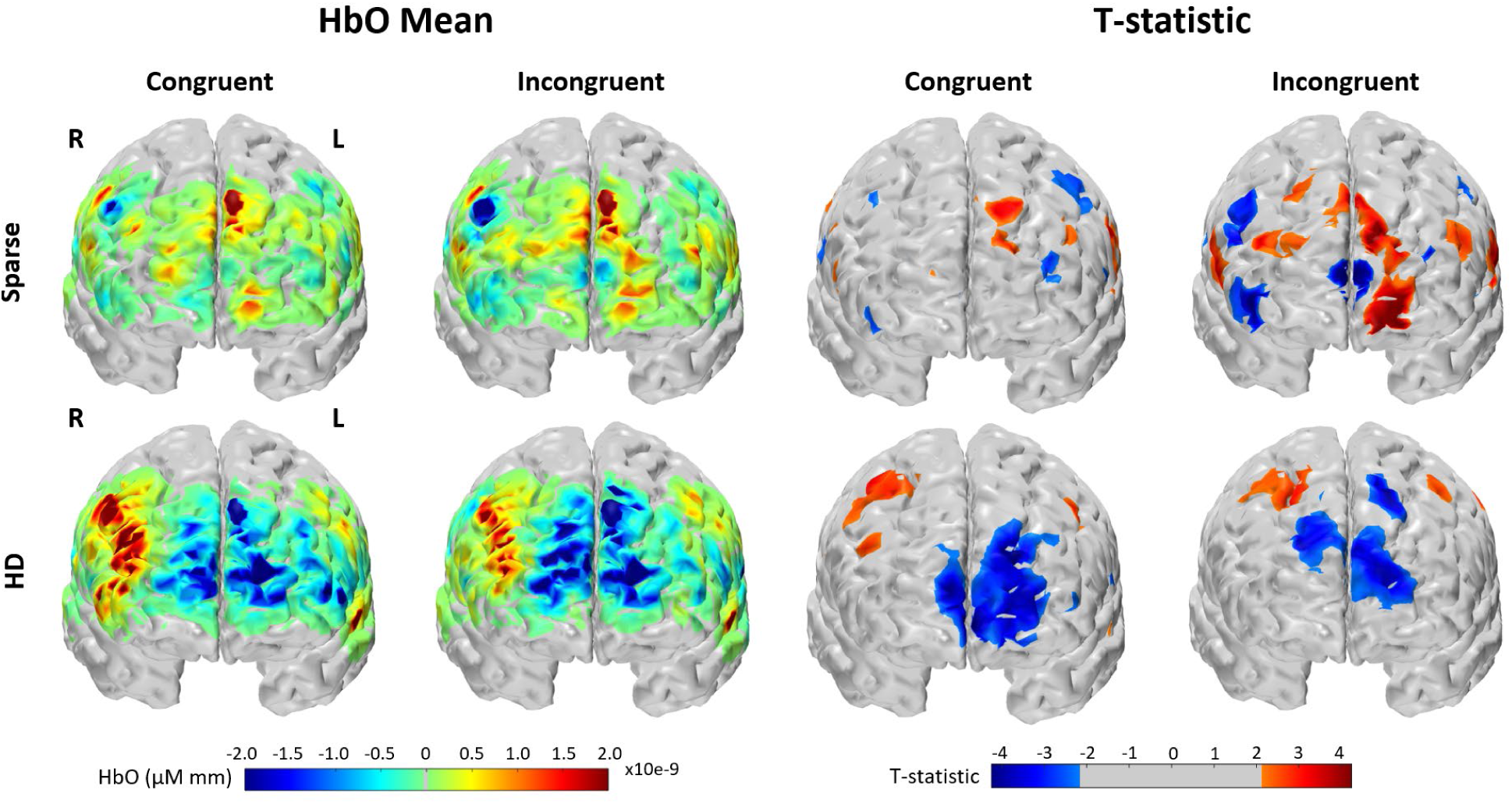
Brain and scalp image space brain response recorded by Sparse and HD arrays during WCS, from Anterior view. “HbO Mean”: Group-average hemodynamic response (HbO) for each condition. “T-statistic”: Group-averaged t-statistic of each vertex is plotted. Color-scale is grey for absolute values less than t-crit = 2.12 as calculated for 17 subjects.

Supplementary materials include similar plots for HbR in channel and brain and scalp image space (Figures S8 and S9), and both HbO and HbR group-average channel timeseries (Figure S10). We have also included all individual subject HbO mean and statistical results for congruent WCS in channel and image space in Supplementary material (Figures S11-S14).

### 3.4 Statistical Comparison

From the ROIs, maximum HbO t-statistics per subject were compared between arrays as in Figure 6. The corresponding average time course from each channel or vertices selected for a given condition, array, and ROI across subjects is shown underneath the bars with which it is associated. (The results for HbR data are in Supplementary Figure S15, and values for both Figure 6 and S15 are provided in Supplementary Table S3.) Of the HD channels selected from each subjects’ ROI for its maximum HbO t-statistic during incongruent WCS, 8 out of the 17 were 19mm. During congruent WCS, 8 of 17 channels selected in the left ROI and 5 of 17 in the right ROI were 19mm. Further comparison of channel statistics if selecting from only the 33mm channels or from only the 19mm channels of the HD array is available in Supplementary material Figure S16 and Table S4.

**Figure 6:**
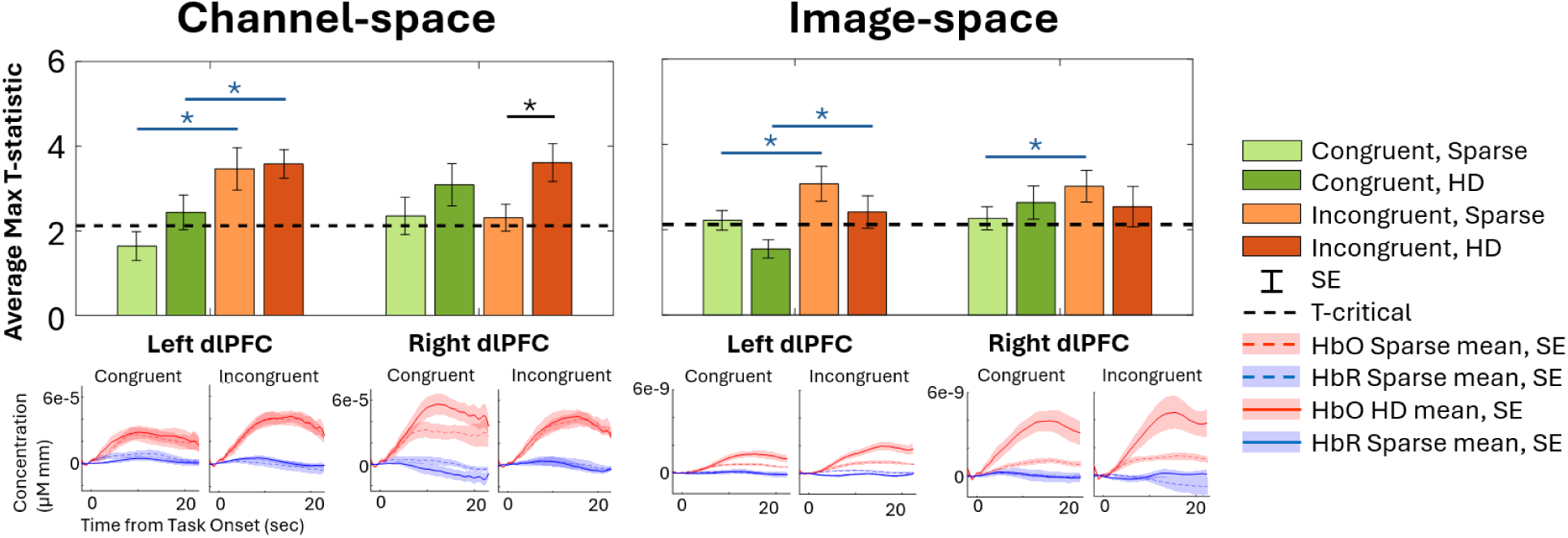
From within the ROIs, group-averaged HbO maximum t-statistics are presented in both channel and brain and scalp image space for each array and WCS conditions. T-critical is 2.12, as calculated for 17 subjects with two-tailed α=0.05. Asterisk indicates p < 0.05 for paired Student’s t-test between arrays (black) and conditions (blue). The subjects’ selected channel or averaged 25 vertices’ concentration time courses are averaged for the timeseries plots. Numerical average, standard error, and paired Student’s t-test values available in Table S3.

The channel statistical data from the HD array was significantly greater in magnitude than that of the Sparse array in the right ROI for both HbO incongruent WCS data and HbR congruent WCS data. In channel space, the HD array consistently provided a higher average maximum t-statistic than did the Sparse array for both HbO and HbR. As a whole, in both channel and image space the HbO statistics from the incongruent condition were greater than those from the congruent condition, with the exception of the right ROI for Sparse in channel space and for HD in image space. The difference was significant in the left ROI for the HbO data in either cap (channel Sparse p = 0.0011, channel HD p = 0.0441; image Sparse p = 0.0306, image HD p = 0.0228; image Sparse right p = 0.0252. All p-values provided in Supplementary Table S3). The average HbO channel and image space statistics also all exceeded the critical t-value, indicating significant brain activation, except the left congruent data for Sparse channel space and HD image space.

## 4 DISCUSSION

Though the prevalence of fiberless HD-DOT systems is increasing^41,70,71^, there has been a lack of adequate characterization of expected improvement of data obtained by such arrays over the commonly available traditional, grid Sparse fNIRS array. To our knowledge, this is the first study to perform a direct statistical comparison of two such arrays in a group of healthy adult subjects, providing statistical analysis in both channel and image space, including short-separation channels in the arrays to perform superficial tissue regression accordingly, and offering comparison for similar task conditions with differing cognitive load.

### 4.1 Channel and Image Statistics Array Comparison

Our analysis quantitatively compared the strength and consistency of signal detected from each array. In channel space, the group-averaged t-statistics from each ROI’s chosen channel showed that the HD array can more consistently capture strength of activation than the Sparse array during both WCS conditions. Similar patterns emerged when looking at the HbR statistics (Figure S15). When comparing t-statistics from each subjects’ ROI vertices from reconstructed images, it was less clear that one array outperforms the other for either condition (Figure 6), though the time courses from selected top vertices show the data analyzed with our image reconstruction pipeline results in higher magnitude of concentration with the HD array.

Interestingly, the 19mm channels comprised approximately half of the HD channels selected from each subjects’ ROI for its maximum HbO t-statistic during incongruent WCS, as well as half of the selected channels for congruent WCS in the left ROI. We anticipated the 33mm channels would yield signal with higher t-statistics than the 19mm channels due to higher brain sensitivity. Previous studies investigated the brain sensitivity of source-detector separations (SDS) ranging from 20 mm to 45 mm and found that for every 10 mm increase in SDS within this range, sensitivity to gray matter increased by approximately 4%. Additionally, the findings indicate that a minimum SDS of 25 mm is required to achieve relative sensitivity greater than 1% at a depth of 11.2 mm into the brain^72^. However, our data suggests that 19mm channels may perform comparatively well to 33mm channels for measuring dlPFC in our healthy adult population, based on the t-statistic metric. For possible explanation as to why the t-statistic is higher, we look at Table 1’s report that the average raw signal SNR of un-pruned 19mm channels (57.3±18.6) was significantly higher than that of the un-pruned 33mm channels (40.9±13.5). Additionally, looking further into the standard error of HbO concentration across blocks, we found that the average SE of the 33mm channels selected for maximum t-statistic was consistently higher than that of the 19mm channels selected for maximum t-statistic (during congruent WCS: 33mm SE is 1.13 ± 0.24 in left ROI and 1.76 ± 0.27 in right ROI, vs the 19mm SE is 0.99 ± 0.18 in left ROI and 1.32 ± 0.25 in right ROI. During incongruent WCS: 33mm SE is 1.09 ± 0.16 in left ROI and 1.37 ± 0.24 in right ROI, vs the 19mm SE is 0.98 ± 0.11 in left ROI and 1.11 ± 0.22 in right ROI).

Our observation of the 19mm’s surprisingly good performance inspired the re-running of the channel space statistical comparison, altered to select a HD channel from either only the ROI’s 19mm channels, or only the ROI’s 33mm channels (Figure S16). We recall that per ROI, there are 6×30mm channels from the Sparse array, 11×19mm channels from the HD array, and 14×33mm channels from the HD array. We found that having both 19mm and 33mm channels consistently yielded the strongest improvement over Sparse data, and that selecting from only 33mm channels did consistently provide slightly higher average maximum t-statistics than when selecting from only 19mm channels. It is not entirely clear if this may have occurred due to having more 33mm channels available than 19mm channels, or from improved signal from the 33mm channels.

We initially also had different expectations for the image space results – specifically, that the HbO pattern from both arrays’ measurements would be similar to that of the channel space statistical comparison in Figure 6 where there is dlPFC activation and mFPC deactivation for both tasks, more so during incongruent WCS. While the HD array resulting pattern of activation aligned with the channel space results, the Sparse array images are almost opposite wherein there is more activation in mPFC and more deactivation in dlPFC.

We explored another method for pushing physiological and measurement noise to the scalp surface while isolating the cortical activity to the brain surface; Gao *et al.* developed a method that uses spatial basis functions modelled as Gaussian kernels with *σ* = 5mm for the brain and *σ* = 20mm for the scalp^27^. This has the advantage of reducing the degrees of freedom of the model and smoothing the resulting images. When we implemented this method, using parameters lambda1 = 0.01 and lambda2 = 0.1 in the sensitivity matrix inversion calculation, it seemed to appropriately handle the HD array data, however, we saw even more dramatic reduction of activation in the Sparse array due to its lack of overlapping channels. This suggests that the traditional Sparse array is ill-posed by nature for reconstructing an appropriate image which separates brain and scalp data.

We suspect that the Sparse array is better represented by not including scalp surface via spatially variant regularization when performing image reconstruction, and we have therefore generated such group result images in Supplementary Figure S17 for the Sparse array which we refer to as “brain only” image reconstruction. In doing so we observe in the images and time course plots that there is strong cross-talk between the HbR and HbO data, which is not present in the regularized data. This demonstrates that the brain only method for reconstructing Sparse data is also unfit. Therefore, for the purpose of directly comparing HD and Sparse data, in our presented image results we chose to apply the same method to both arrays and include that in analysis in Figure 6. Because our analysis selects the top vertices based on the HbO increase in the ROIs (and HbR decrease in the ROIs as shown in the supplementary material) we can still compare potentially-relevant results, but we argue that it may be inappropriate to assess Sparse image results as a whole.

For both arrays, it is possible that our ability to perform a more robust statistical comparison of image space results was hindered by the limitation that, like many other fNIRS studies, we did not have subject-specific MRIs to support more accurate probe localization. Rather, we used the AtlasViewer-provided Colin27 head atlas across all subjects.

### 4.2 Capturing Different Cognitive Load

This study allowed us to observe each arrays’ comparative performance for different cognitive loads. The Stroop effect was clearly demonstrated in our data by the increased activation during incongruent task as compared to congruent in both caps (activation pattern is discussed further in section 4.3) and performance metrics were comparable to what was expected based on previous literature^53,73^, so we are confident that our study setup generated desired brain activity. The performance metrics’ lack of significant difference in accuracy or response time between Sparse and HD array verifies comparability of the WCS brain response between arrays. The presence of significantly longer response time during incongruent WCS than during congruent WCS while maintaining accuracy, both when wearing Sparse and HD array, suggests and supports there is a higher cognitive load when performing incongruent WCS (*Table 2*). This is expected since the congruent condition does not require interference resolution and response inhibition to identify the correct response, which supplies non-interfering information.

We found that both the Sparse and HD arrays’ channel and image statistical data in the left ROI were significantly different between congruent and incongruent WCS. Other studies have also found the left PFC to be more involved than the right in the Stroop effect^74^. While this might indicate either array is suitable to find difference between task-specific activation, looking at the activation patterns of each condition independently yields a more nuanced perspective. In channel space for the incongruent task, both arrays captured the presence of activation as seen in the statistical visualization of Figure 4, to varying degrees of localization discussed further in section 4.3. However, there was a markedly different ability between the arrays to capture activation during the congruent task. The HD array still captured robust activation patterns in channel space; by contrast the Sparse array results had very few channels – none in the left ROI – that detect significant activation at the group level. This aligns with the channel statistical comparison in Figure 6, in which we observe that for congruent WCS in the left ROI the averaged maximum t-statistic does not exceed the critical t-value. Looking at image results in Figure 5, the Sparse array is similarly devoid of activation in the ROIs. For both tasks, though the statistics in Figure 6 would suggest HD does not outperform Sparse especially in the left ROI, the visualization in Figure 5 clearly indicates the HD array consolidates, or localizes, activation for both tasks.

This has implications for future array choice which should depend on a given paradigm’s difficulty or expected magnitude of activation, and intention to analyze results in channel or image space. It is worth noting that most of our subjects have similarly high educational attainment, and the age range is limited which may skew the data. Future work might implement a similar comparative method with other tasks and conditions and across other populations to build a bank of array detection characterizations. This could enable better-informed selection of paradigm pairing with selected array density and location, whether for research, clinical use, or other purposes. Additionally, due to the importance of differentiating subject-specific activity between conditions^75^ or condition-specific activity between subjects^3,7,52^, more work could look at other condition-specific metrics, such as signal latency, integral value, or centroid value^76,77^.

### 4.3 Localization Comparison

The pattern of HbO activation during WCS as detected by both arrays in channel space and by HD array in image space matched that of previous fNIRS and fMRI findings specific to WCS^50–52,54,73^ – that is, lateral activation and medial deactivation, more strongly in the incongruent condition than in the congruent condition (Figure 4, 5). From the visualization of both arrays in channel and HD array in image space (Figure 4, 5) as well as the statistical comparison in channel space (Figure 6), the incongruent task seemed to elicit a more uniformly bilateral response than the congruent task for which there is more activation present in the right ROI than the left. This hemispherical activation difference for congruent WCS was not observed in most other studies^50,53,54^. However, it is known that the left dlPFC is implicated in interference processes like that induced by Stroop effect^56,57^, so it follows that the difference of activation between congruent and incongruent tasks is greater in the left than right dlPFC which was true of our data as well (Figure 6). Though we did not expect the lateral difference for congruent data, there is another published work of healthy adults which also found lesser activation in the left PFC than the right PFC for their non-incongruent task^55^.

Looking at the visualized regions of significance-thresholded activity (Figures 4 and 5), especially for the higher-cognitive load incongruent task, it becomes clear that the HD array better localizes the activation in both channel and image space. This validates the Monte Carlo simulated photon migration results which showed reduced localization error and improved sensitivity in the HD array (Figures 1 and S1). This also supports previous literature naming the same advantage in infant populations and without applying short-separation regression to all the data^31,49^. This is most dramatically illustrated in the reconstructed image data (Figure 5), for which the active region detected by HD is continuous and contains a relatively centered focus of higher activity. By contrast, the active region detected by Sparse is discontinuous and lacks a centered focus of activity. Our study, therefore, translates some of existing findings to the 3cm grid array which is commonly available and in use and to an adult population, better enabling comparison of the systems and for broader application.

### 4.4 Signal Quality and Data Collection

Since our comparison occurred over the PFC, the probe designs did not cover as much hair as they would have for other regions like parietal or occipital. This enabled us to perform the study without the need to assess how the presence of hair might affect the layouts differently, providing more of an ideal baseline comparison between the two. The left and right extremities of the arrays, though, do extend into hair regions especially on the superior boundaries of the cap, and we would predict that the HD array signal is more impacted by hair. Surprisingly, we saw that the SNR of the Sparse and HD array were not significantly different for non-pruned channels (p=0.36), which demonstrates promise for comparable signal quality. However, the significantly greater percentage and number of channels pruned for low SNR and low raw intensity from the HD array than for Sparse point to increased difficulty in achieving optimized signal via the HD array. Even so, we note that on average we still had over three times as many channels available from the HD array as compared to Sparse; within our ROI we had over four times as many. On average, we retained 91.6% of channels in the HD array which is higher than the channel retention for previous HD studies^31^. The HD pruned channels were mainly located in the superior portions of the array so we conclude the increased proportion of pruned channels from the HD array was due to presence of hair (Figure S19). Future work may characterize these metric comparisons in other regions of the head where there is more hair and hopefully elaborate on methods to reduce the percentage of HD channels pruned for SNR and low signal. We do recommend that users of fNIRS systems follow our described methods of cap placement and scalp coupling to enhance signal quality and use of the ninjaCap^60^ itself, in combination with other documented techniques^78–82^, and look forward to further developments of user techniques or technologies to ensure good signal quality optimization across demographics. Anecdotally, we can speak to our subjects having a range of hair color, shape, and texture, as well as a range of skin tone, thus verifying the applicability of the findings across various racial demographics. However, we recognize the absence in our study and suggest future studies to regularly collect metrics of hair color, shape, and texture as well as skin melanin pigmentation metrics to continue documenting and improving fNIRS ability to accommodate for a full range of demographics^82^.

A challenge we faced was that the number of optodes we had available allowed for only one array to be assembled at a given time. Subjects provided their estimated head circumference ahead of the session, and therefore we could populate optodes on the NinjaCap for either HD or Sparse for the estimated size. Unfortunately, subjects did not always provide an accurate head measurement as determined once we measured after consenting (differences were no larger than 2cm). Due to time constraints for the full session duration, we chose to proceed with the pre-assembled cap size; when moving optodes to the other layout for the second run of the session we used the same cap size for per-subject size consistency. If a cap was too large for the subjects head it may loosen the optode contact with the subjects’ head, which we accounted for by creating a fold in the occipital region (similar to a sewing dart) and securing with a clip. While using caps of incorrect head size would not affect pairwise comparison of data, it is likely to, in small ways, have affected localization of group results^83^. We recommend future work to navigate a challenge like this by a variety of methods, such as pre-study head measurements, digitizing probe localization after cap placement, using a size-adjustable cap design, or simply populating multiple cap sizes if the resources are available.

In order to perform a direct comparison of our HD array to the Sparse grid array, we were constrained to using two separate printed caps and populating them separately due to non-overlapping optode locations. Though this potentially introduced variability in cap placement per subject, our pair-wise analytical methods overcome this by selecting one channel or vertex per ROI with greatest t-statistic. Additionally, any effect of variability in per-subject cap placement in group results would not be any greater than effects due to inter-subject cap placement variability.

## 5 CONCLUSION

As fNIRS development and research progresses toward whole-head HD systems, this work’s characterization in the PFC of the improvement our HD layout affords over that of the traditional sparse array may apply across the whole-head. Several key implications of Sparse and HD array use emerged from our study which can inform future selection of array and paradigm to best elicit and capture functional activity. Comparison of congruent task results suggest that the Sparse array is inadequate for capturing brain activity when used for tasks which have a lower cognitive load in healthy adult subjects; conversely, our HD array is sufficient for capturing such brain activity. For both WCS conditions, localization is better achieved with the HD array and image reconstruction can be more appropriately performed with the HD array. If localization is not of high importance and analysis of image space results is not necessary, our incongruent results indicate that either array can be sufficient to capture the presence of activation when designed with continuous coverage over the ROI. The WCS task is relevant for many studies in psychiatric well-being and other cognitive applications, thus these results can hopefully translate directly to inform future studies using WCS and related Stroop or response-inhibition paradigms. We hope our methods and findings offer a foundation on which ongoing fNIRS array comparisons, expansions, and applications may build.

## 6 DISCLOSURES

The authors declare that there are no financial interests, commercial affiliations, or other potential conflicts of interest that could have influenced the objectivity of this research or the writing of this paper.

## 7 CODE AND DATA AVAILABILITY

Both the Sparse and HD frontotemporal array designs are openly accessible at https://openfnirs.org/hardware/ninjacap/ in file formats of.SD and.SNIRF for use with the AtlasViewer and Homer3 platforms, and in three circumference size files (54cm, 56cm, 58cm) in.stl and Cura file formats for 3D printing. Code to run the Word-Color Stroop paradigm in PsychoPy is publicly accessible at https://github.com/andersonjessie/WordColorStroop). De-identified recorded fNIRS and task performance data are available in separate datasets at openneuro.org.

## 8 AUTHOR CONTRIBUTIONS

JEA, DAB, and MAY conceptualized the research question and analysis. JEA designed the experimental approach and protocol and executed the experiments. JEA, SK, and WJO prepared the materials for data collection. JEA, PR, and PF managed the recruitment of participants. JEA, MAY, DAB, and LC analyzed and discussed the data. JEA and LC drafted the original manuscript. All authors reviewed, provided feedback, and edited the manuscript.

## 9 ACKNOWLEDGMENTS

We thank Yuanyuan Gao, Antonio Ortega, Sudan Duwadi, Darash Desai, Alexander Von Lühmann, Jack Giblin, Xiaojun Cheng, Byungchan (Kenny) Kim, Chantal Stern, Alice Cronin-Golomb, Rini Kaplan, and Neila Gross for helpful lab training and insightful discussions and feedback. This project was supported by NIH NEW grant U01-EB 029856 and Boston University research funds.

## 10 AUTHOR BIOGRAPHIES

Jessica E. Anderson is a PhD candidate in Biomedical Engineering at Boston University. Her work focuses on fNIRS’ development, application to studying emotional regulation in developing populations, and translation to global contexts.

Laura Carlton a PhD candidate in Biomedical Engineering at Boston University. She completed her undergraduate degree in bioengineering at McGill University. Her current research centers on developing advanced computational methods and data analysis techniques to enhance the application of fNIRS technology in naturalistic studies, with the aim of improving its utility in real-world environments.

Sreekanth Kura is a Research Fellow at Boston University, specializing in fNIRS system development and acquisition software. His work also encompasses computational methods for optical microscopy.

W. Joseph O’Brien, MS, works primarily with the design development of optical neuroimaging equipment. While with the Boas Lab, he worked towards the design of high-density fNIRS (HD-fNIRS) and the integration of other modalities into a whole head HD-fNIRS capable device.

De’Ja Rogers is a biomedical engineering doctoral research fellow at Boas Lab, Boston University. She received her BS degree in electrical and electronics engineering from Norfolk State University and her MS degree in biomedical engineering from Boston University. She focuses on optimizing the combination functional near-infrared spectroscopy (fNIRS) and electroencephalography (EEG), with the goal of investigating neurodegeneration in the future.

Parisa Haji Rahimi holds degrees in Physics, and Business Administration, with a focus on artificial intelligence and deep learning. She is passionate about leveraging these technologies to solve real-world problems across various industries.

Parya Y. Farzam is a research fellow at the Boston University (BU) Neurophotonics Center. She uses fNIRS to perform cognitive neuroscience study and mainly focuses on the brain function of children with autism spectrum disorder. She prior to joining BU, she was a visiting student at the Martinos Center for Biomedical Imaging at Massachusetts General Hospital. She has a bachelor’s degree in software engineering. She has hands-on experience with fNIRS in the research field of cognitive neuroscience, and also using diffuse correlation spectroscopy (DCS) on patients with stroke.

Muhammad H. Zaman, PhD, is an HHMI professor of Biomedical Engineering and Global Health at Boston University and the Director of Center on Forced Displacement. His research focuses on disease dynamics and access to healthcare among forcibly displaced persons - including refugees, internally displaced persons and stateless communities.

David A. Boas, PhD, is a professor of biomedical engineering at Boston University. He is the founding president of the Society for Functional Near-Infrared Spectroscopy and founding Editor-in-Chief of the SPIE journal Neurophotonics. He received the Britton Chance Biomedical Optics Award in 2016 for his development of several novel, high-impact biomedical optical technologies in the neurosciences, as well as following through with impactful application studies, and fostering the widespread adoption of these technologies.

Meryem A. Yücel, PhD, is a research associate professor at Boston University (BU). Prior to her position at BU, she was an assistant in biomedical engineering at Massachusetts General Hospital and an instructor at Harvard Medical School, Radiology. Her primary research interest is to understand how the brain works in health and disease. Throughout her career, she has gained expertise in mathematical modeling of biological systems and functional brain imaging (fNIRS, fMRI, and EEG).

## Notes

### Competing Interest Statement

The authors have declared no competing interest.

