## Supplementary for "High-Density Multi-Distance fNIRS Enhances Detection of Brain Activity during a Word-Color Stroop Task"

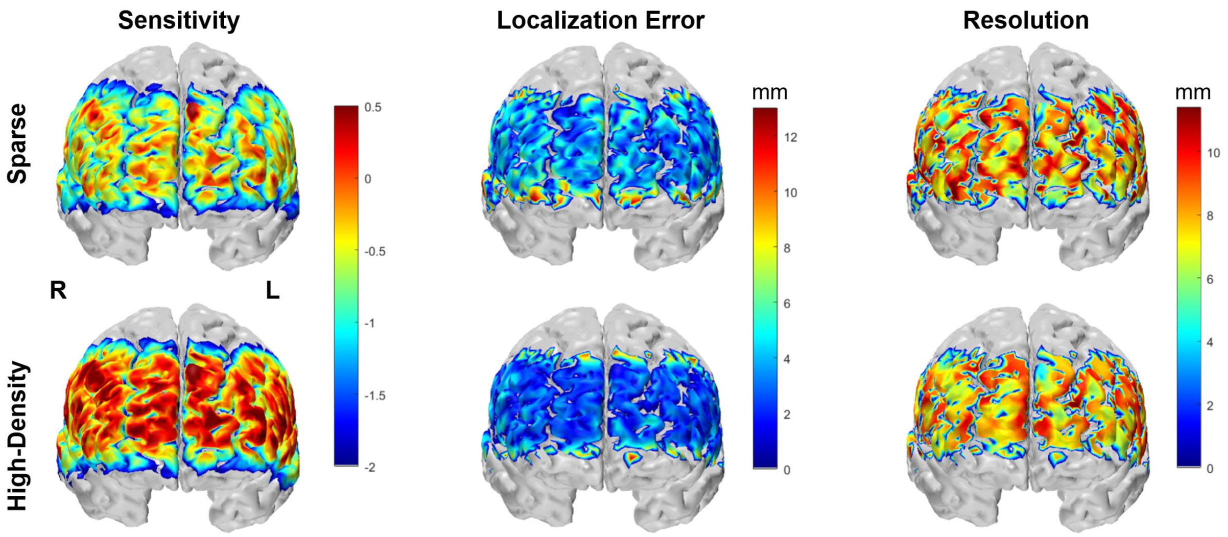


Figure S7: Sensitivity, localization error and resolution maps are shown for sparse (A) and high-density (B) optode arrays. Sensitivity profile is on a log 10 scale; vertices with values > 0.01 are not masked and not considered part of the relevant sensitivity profile.

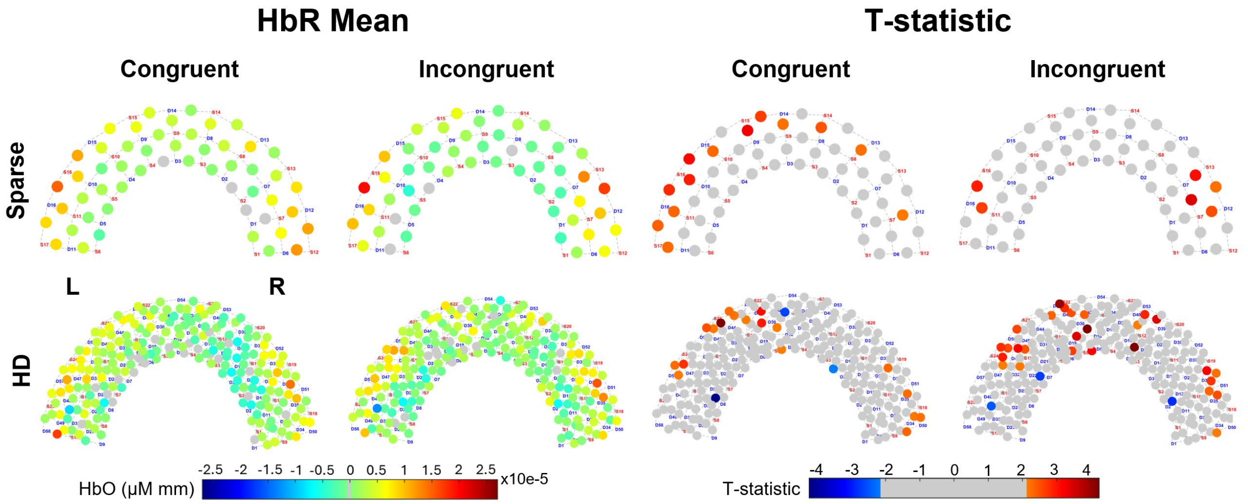


Figure S8: Channel-level brain response recorded by Sparse and HD arrays during WCS. “HbR Mean”: Group-average hemodynamic response (HbR) for each channel, averaged across 7 to 18sec of the blocks for each condition. “T-statistic”: Group-averaged t-statistic of each channel is plotted. Color-scale is grey for absolute values less than two-tailed t-critical = 2.12 as calculated for 17 subjects.

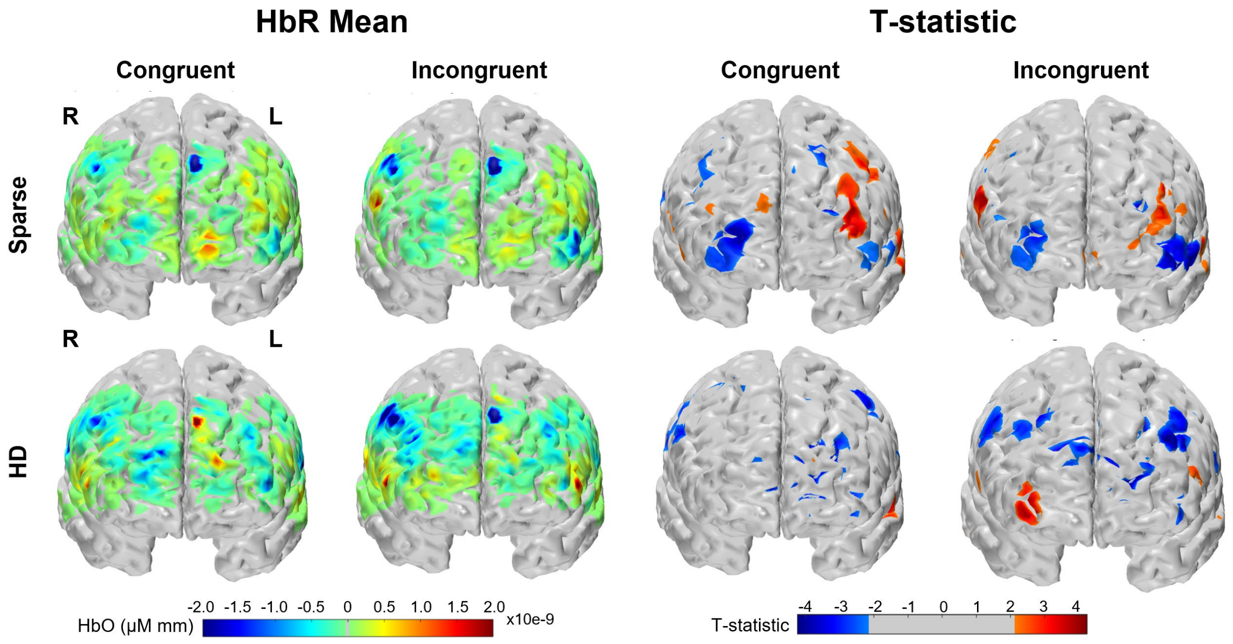


Figure S9: Brain and scalp image space brain response recorded by Sparse and HD arrays during WCS, from Anterior view. "HbR Mean": Group-average hemodynamic response (HbR) for each condition. "T-statistic": Group-averaged t-statistic of each vertex is plotted. Color-scale is grey for absolute values less than two-tailed t-critical = 2.12 as calculated for 17 subjects.

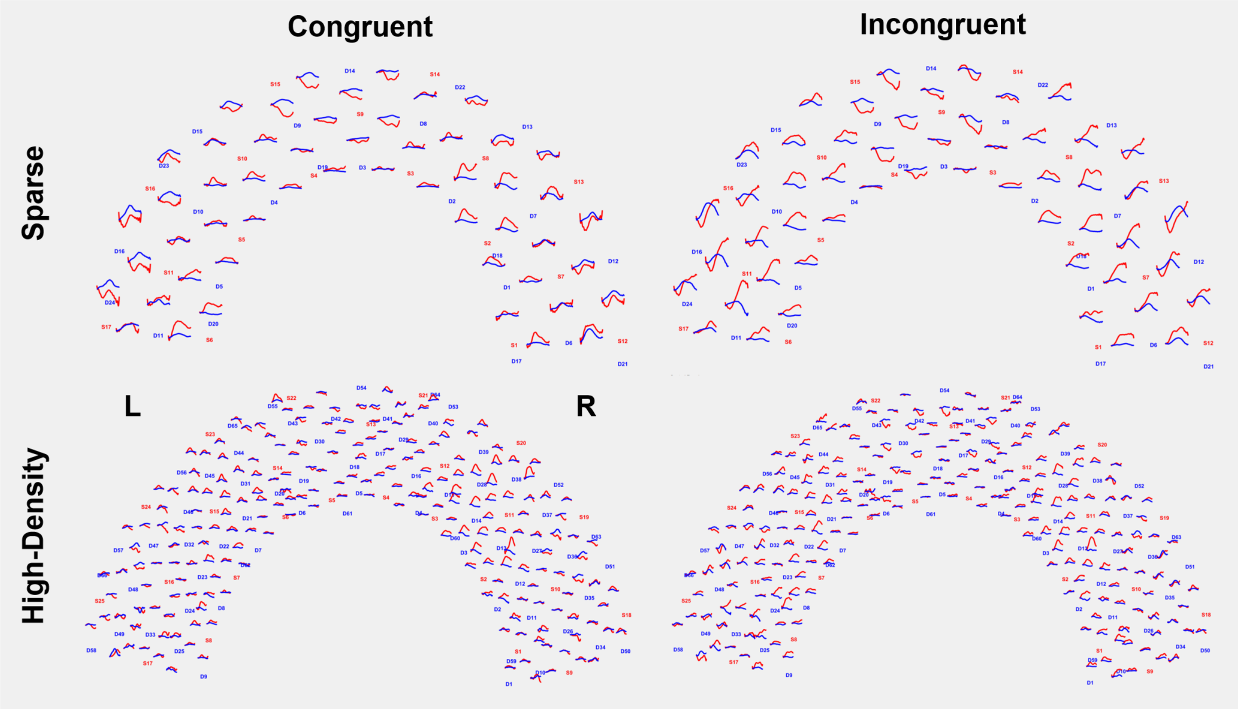


Figure S10: Group-average hemodynamic response (HbO and HbR) from -2 to 25sec for each channel. Red lines indicate HbO, blue lines indicate HbR. Y-axes agree within a result (i.e. Sparse Congruent) but are not aligned across results.

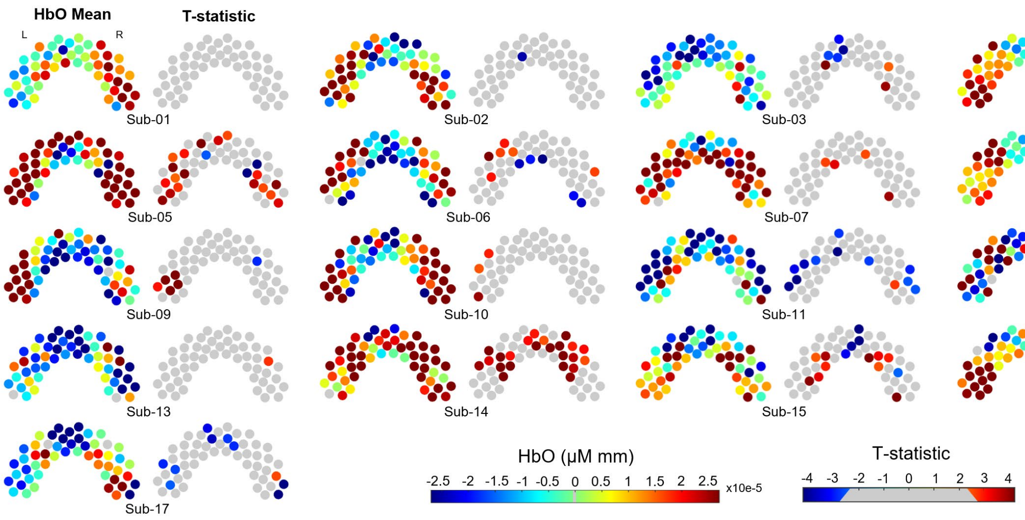


Figure S11: Channel space brain response recorded by Sparse array during Incongruent WCS, from posterior view. Each subject's HbO mean and t-statistic across blocks is side-by-side. T-statistic color-scale is grey for absolute values less than two-tailed t-critical ranging 2.306 to 2.776 as calculated for each subject's blocks used in analysis (ranges from 5 to 9).

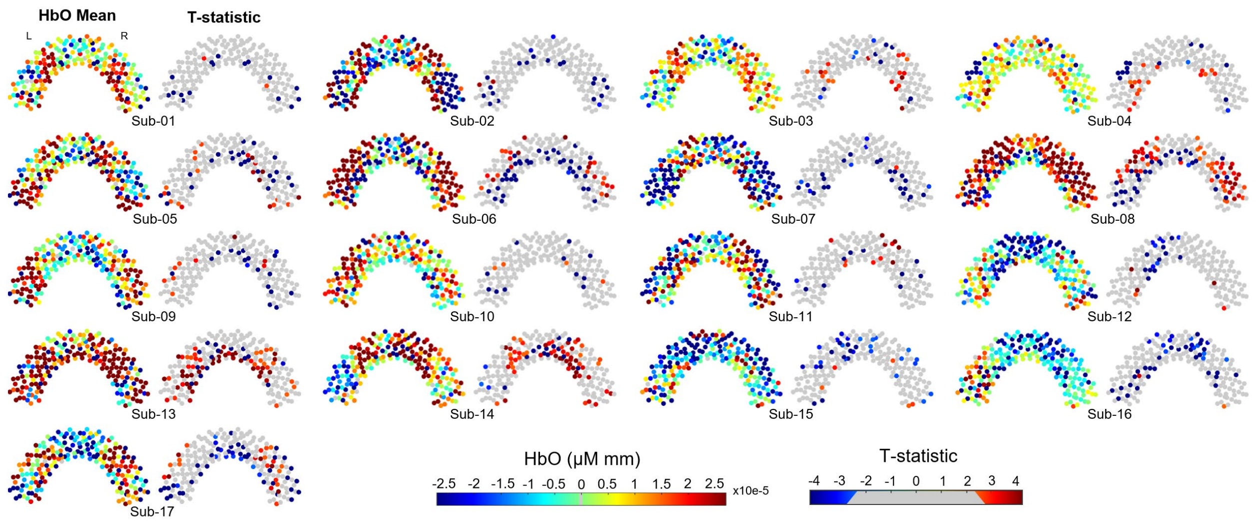


Figure S12: Channel space brain response recorded by HD array during Incongruent WCS, from posterior view. Each subject's HbO mean and t-statistic across blocks is side-by-side. T-statistic color-scale is grey for absolute values less than two-tailed t-critical ranging 2.306 to 2.776 as calculated for each subject's blocks used in analysis (ranges from 5 to 9).

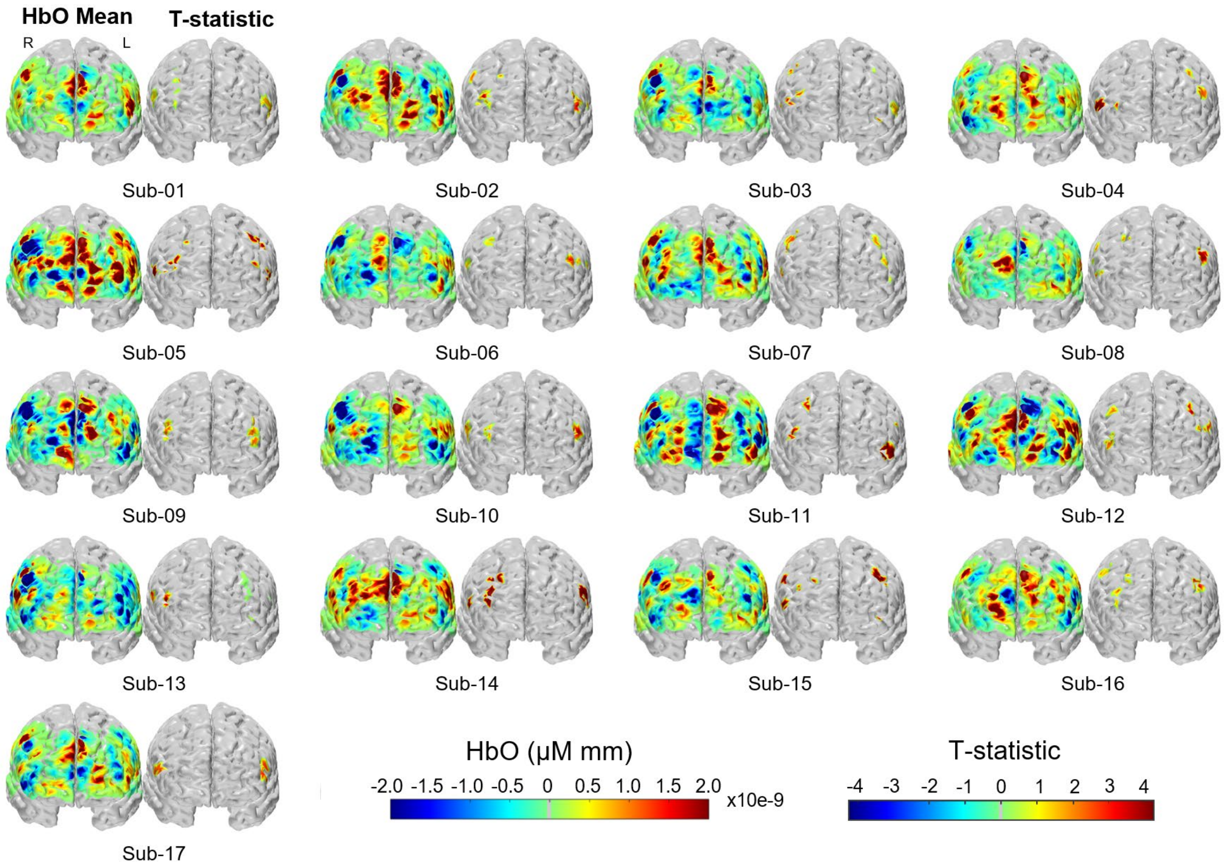


Figure S13: Brain and scalp image space brain response recorded by Sparse array during Incongruent WCS, from anterior view. Each subject's HbO mean and t-statistic across blocks is side-by-side. The t-statistic images display the top 25 vertices per ROI and were used in analysis.

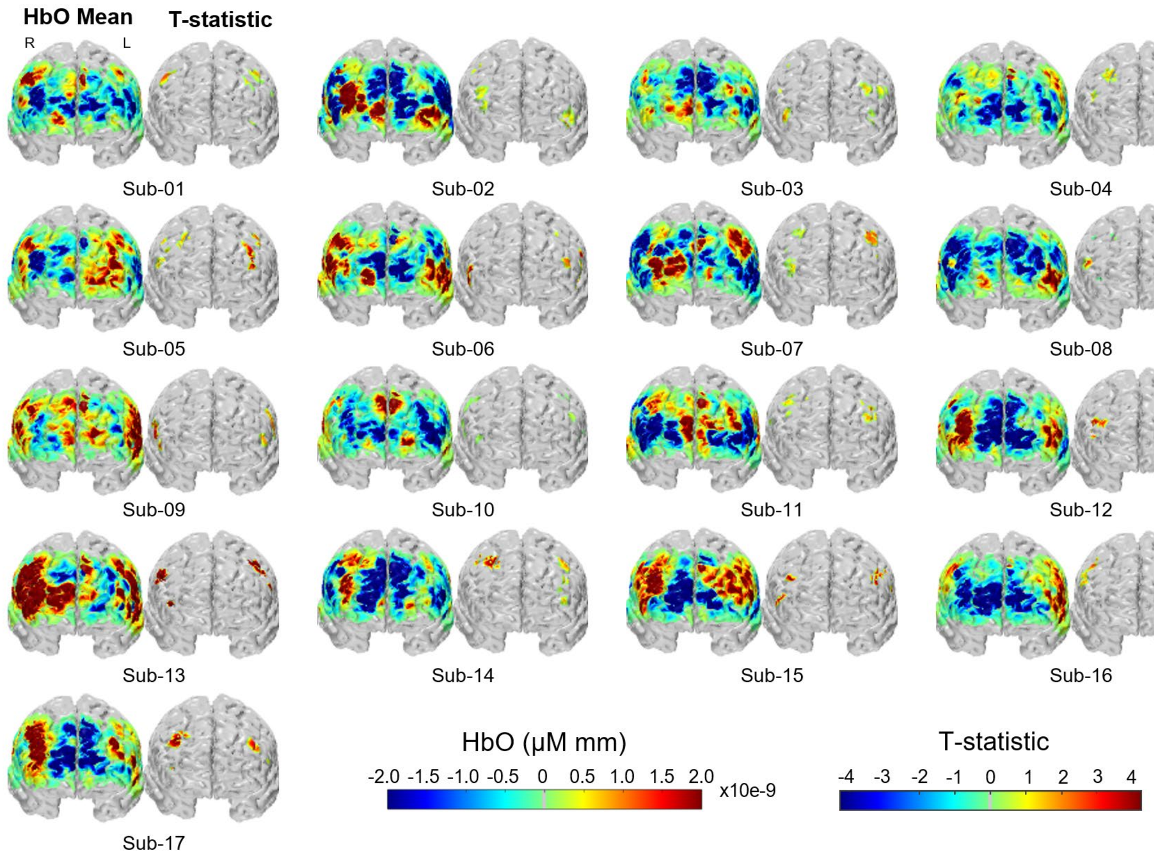


Figure S14: Brain and scalp Image space brain response recorded by HD array during Incongruent WCS, from anterior view. Each subject's HbO mean and t-statistic across blocks is side-by-side. The t-statistic images display the top 25 vertices per ROI and were used in analysis.

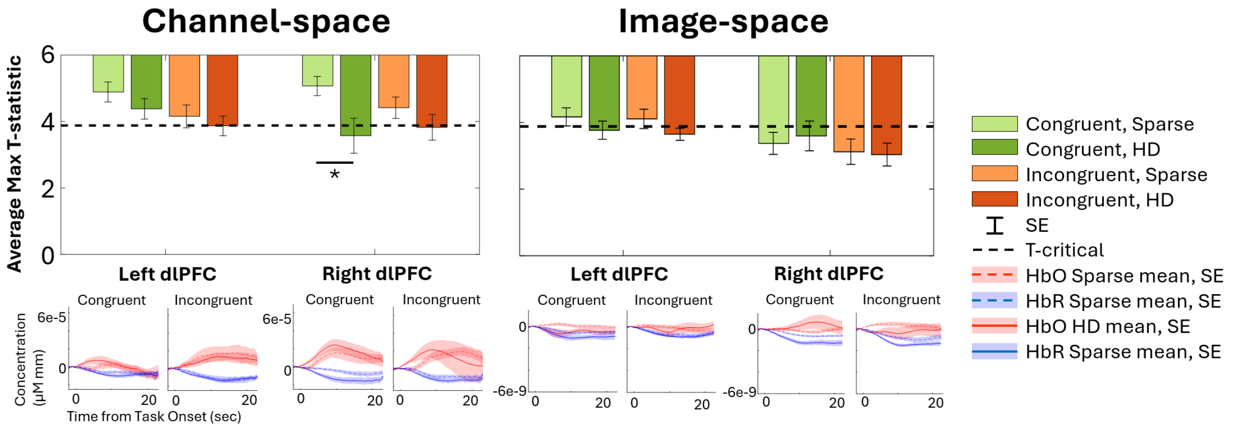


Figure S15: From within the ROIs, group-averaged HbR minimum t-statistics are presented in both channel and brain and scalp image space for each array and WCS conditions. T-critical is 2.12, as calculated for 17 subjects with two-tailed α=0.05. Asterisk indicates $p < 0.05$ for paired Student's t-test between arrays (black) and conditions (blue). The subjects' selected channel or averaged 25 vertices' concentration time courses are averaged for the timeseries plots. Numerical average, standard error, and paired Student's t-test values available in Table S1.

| HbO | Channel-Space | | | | | | Brain and Scalp Image-Space | | | | | |
|---|---|---|---|---|---|---|---|---|---|---|---|---|
| | Congruent | | Incongruent | | p value, Condition | | Congruent | | Incongruent | | p value, Condition | |
| | Left | Right | Left | Right | Left | Right | Left | Right | Left | Right | Left | Right |
| **Sparse** | 1.64 ± 0.34 | 2.35 ± 0.44 | 3.47 ± 0.50 | 2.31 ± 0.32 | *0.0011 | 0.94 | 2.22 ± 0.23 | 2.27 ± 0.27 | 3.08 ± 0.41 | 3.02 ± 0.37 | *0.0306 | *0.0252 |
| **HD** | 2.44 ± 0.41 | 3.09 ± 0.50 | 3.58 ± 0.34 | 3.61 ± 0.45 | *0.0441 | 0.46 | 1.55 ± 0.21 | 2.64 ± 0.39 | 2.42 ± 0.38 | 2.54 ± 0.47 | *0.0228 | 0.81 |
| **p_value, array** | 0.12 | 0.31 | 0.84 | *0.0273 | | | 0.09 | 0.42 | 0.30 | 0.42 | | |

| HbR | Channel-Space | | | | | | Brain and Scalp Image-Space | | | | | |
|---|---|---|---|---|---|---|---|---|---|---|---|---|
| | Congruent | | Incongruent | | p value, Condition | | Congruent | | Incongruent | | p value, Condition | |
| | Left | Right | Left | Right | Left | Right | Left | Right | Left | Right | Left | Right |
| **Sparse** | -1.11 ± 0.30 | -0.93 ± 0.29 | -1.84 ± 0.34 | -1.58 ± 0.32 | 0.11 | 0.10 | -1.84 ± 0.27 | -2.63 ± 0.33 | -1.89 ± 0.29 | -2.88 ± 0.38 | 0.87 | 0.62 |
| **HD** | -1.62 ± 0.31 | -2.43± 0.53 | -2.13 ± 0.29 | -2.17 ± 0.39 | 0.19 | 0.62 | -2.23 ± 0.27 | -2.41 ± 0.44 | -2.35 ± 0.18 | -2.96 ± 0.34 | 0.71 | 0.26 |
| **p_value, array** | 0.15 | *0.0179 | 0.58 | 0.29 | | | 0.27 | 0.70 | 0.20 | 0.86 | | |

Table S3: Corresponds to data presented in Figure 6 and Figure S9. Group-average of each subject's channels' or vertices' maximum t-stat for a given concentration, WCS condition, ROI, and array. Numbers per array are  the average ± standard error. P-value results from paired-ttest between the arrays' maximum t-statistics.

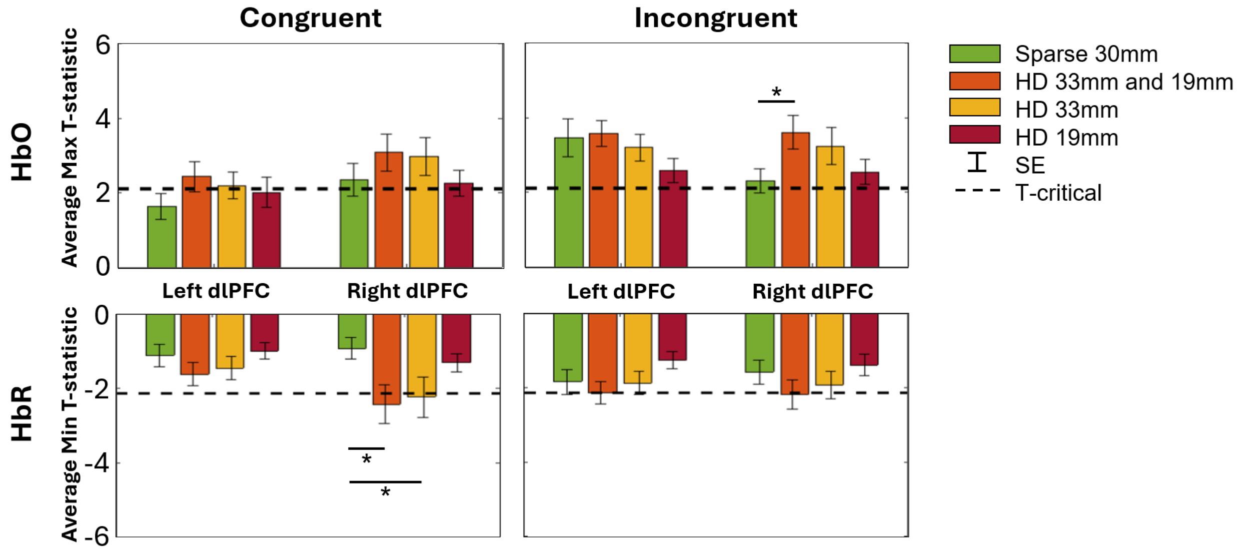


Figure S16: From within the ROIs, group-averaged maximum HbO and minimum HbR t-statistics are presented in channel space for each WCS condition and array channel length groups from which the channels are selected. T-critical is 2.12, as calculated for 17 subjects with two-tailed α=0.05. Asterisk indicates p < 0.05 for paired Student's t-test between arrays. Numerical average, standard error, and paired Student's t-test values available in Table S4.

| **HbO** | **Channel-Space** | | | |
|---|---|---|---|---|
| | Congruent | | Incongruent | |
| | Left | Right | Left | Right |
| **Sparse: 30mm** | 1.64 ± 0.34 | 2.35 ± 0.44 | 3.47 ± 0.50 | 2.31 ± 0.32 |
| **HD: 19 or 33mm** | 2.44 ± 0.41 | 3.09 ± 0.50 | 3.58 ± 0.34 | 3.61 ± 0.45 |
| **p_value** | 0.12 | 0.31 | 0.84 | *0.0273 |
| **HD: 33mm** | 2.21 ± 0.35 | 2.98 ± 0.52 | 3.20 ± 0.36 | 3.25 ± 0.49 |
| **p_value** | 0.21 | 0.40 | 0.68 | 0.10 |
| **HD: 19mm** | 2.03 ± 0.41 | 2.26 ± 0.36 | 2.58 ± 0.33 | 2.55 ± 0.33 |
| **p_value** | 0.45 | 0.89 | 0.11 | 0.62 |

| **HbR** | **Channel-Space** | | | |
|---|---|---|---|---|
| | Congruent | | Incongruent | |
| | Left | Right | Left | Right |
| **Sparse: 30mm** | -1. 11 ± 0.30 | -0.93 ± 0.29 | -1.84 ± 0.34 | -1.58 ± 0.32 |
| **HD: 19 or 33mm** | -1.62 ± 0.31 | -2.43 ± 0.53 | -2.13 ± 0.29 | -2.17 ± 0.39 |
| **p_value** | 0.15 | *0.0179 | 0.58 | 0.29 |
| **HD: 33mm** | -1.45 ± 0.32 | -2.24 ± 0.55 | -1.87 ± 0.31 | -1.93 ± 0.37 |
| **p_value** | 0.334 | *0.0413 | 0.95 | 0.52 |
| **HD: 19mm** | -0.99 ± 0.22 | -1.31 ± 0.24 | -1.25 ± 0.23 | -1.39 ± 0.29 |
| **p_value** | 0.74 | 0.30 | 0.21 | 0.65 |

Table S4: Group-average of each subject's channels' maximum t-stat for a given concentration, WCS condition, ROI, and array. HD array options include when selecting from among all channels in an ROI (whether 33mm or 19mm), from among only 33mm channels, and from among only 19mm channels. Number per array are the average ± standard error, units of μM mm. P-value results from paired-ttest between arrays' maximum t-statistics from the Sparse and each of three HD options. Corresponds to data in Figure S16.

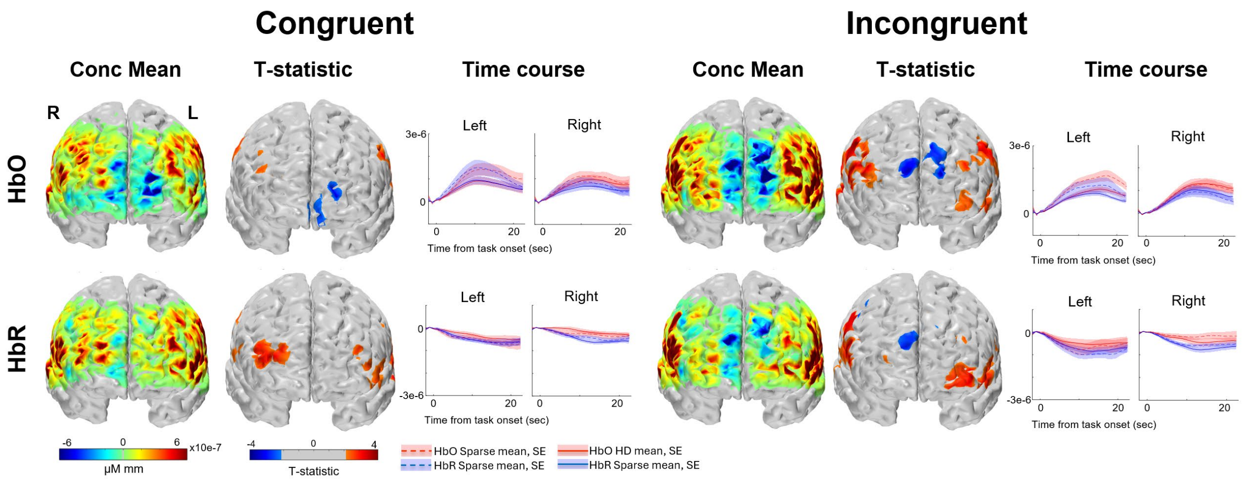


Figure S17: Performing brain only image reconstruction without regularization parameters, the image space response recorded by Sparse arrays during WCS, from Anterior view. “Concentration Mean”: Group-average hemodynamic response for each condition. “T-statistic”: Group-averaged t-statistic of each vertex is plotted. Color-scale is grey for absolute values less than two-tailed t-critical = 2.12 as calculated for 17 subjects. “Time course”: the group average of each subjects’ top 25 vertices by t-statistic ranking.

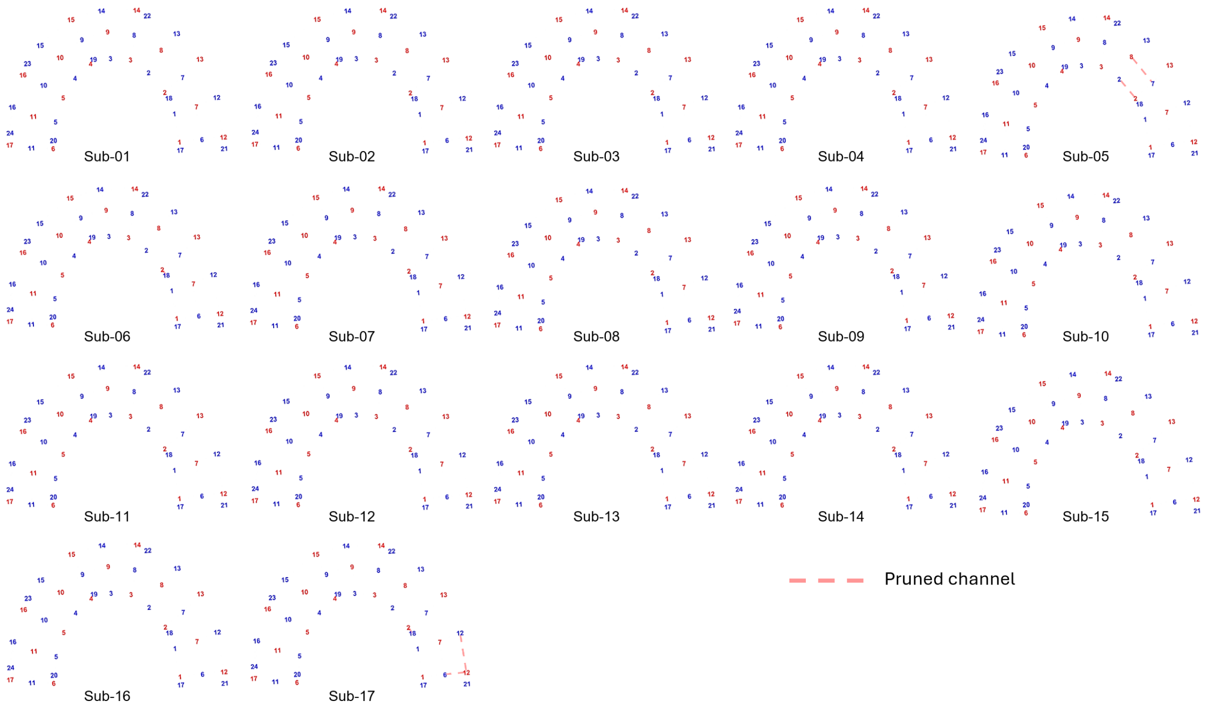


Figure S18: Visualizing subjects' pruned channels for the Sparse array, indicated by red dashed line.

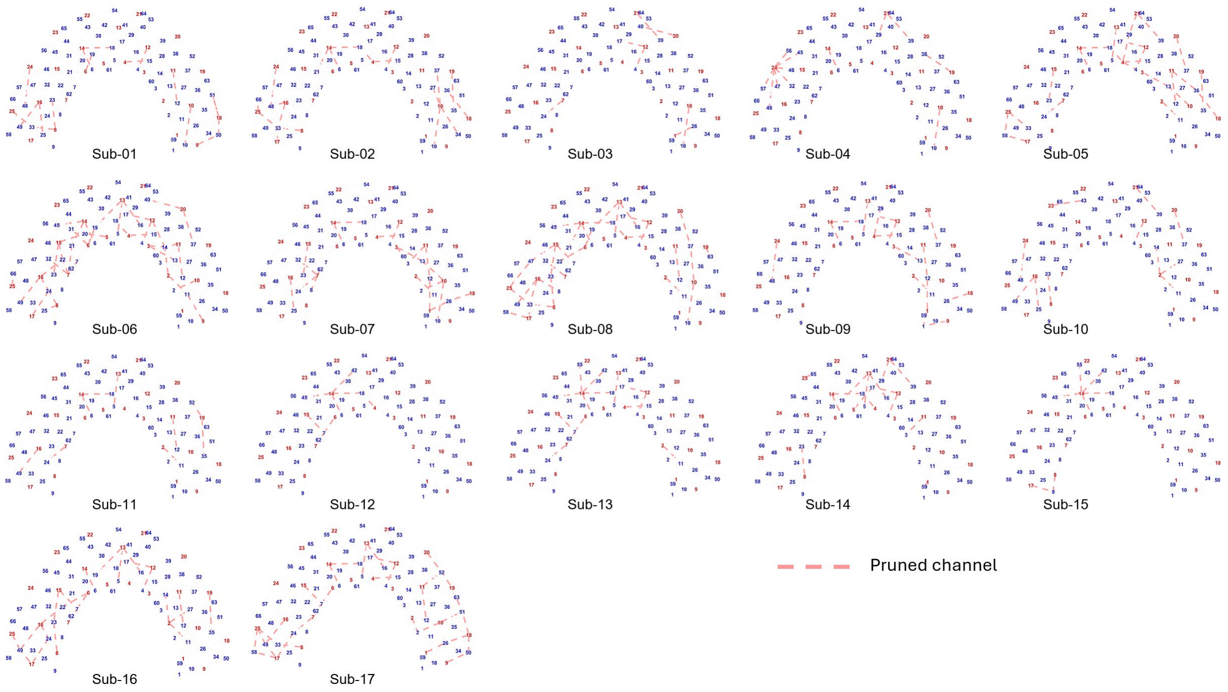


Figure S19: Visualizing subjects' pruned channels for the HD array, indicated by red dashed line.
